# An RNA Splicing System that Excises Transposons from Animal mRNAs

**DOI:** 10.1101/2025.02.14.638102

**Authors:** Long-Wen Zhao, Christopher Nardone, Joao A. Paulo, Stephen J. Elledge, Scott Kennedy

## Abstract

All genomes harbor mobile genetic parasites called transposable elements (TEs). Here we describe a system, which we term SOS splicing, that protects *C. elegans* and human genes from DNA transposon-mediated disruption by excising these TEs from host mRNAs. SOS splicing, which operates independently of the spliceosome, is a pattern recognition system triggered by base-pairing of inverted terminal repeat elements, which are a defining feature of the DNA transposons. We identify three factors required for SOS splicing in both *C. elegans* and human cells; AKAP17A, which binds TE-containing mRNAs; the RNA ligase RTCB; and CAAP1, which bridges RTCB and AKAP17A, allowing RTCB to ligate mRNA fragments generated by TE excision. We propose that SOS splicing is a novel, conserved, and RNA structure-directed mode of mRNA splicing and that one function of SOS splicing is to genetically buffer animals from the deleterious effects of TE-mediated gene perturbation.

## Main

TEs that integrate into or near genes often disrupt gene function. To limit such disruptions, organisms possess epigenetic systems that inhibit TE expression and replication^1–4^. Despite these systems, TEs constitute a substantial (3-80%) fraction of extant genomes^5^, indicating that TE-silencing systems are not full-proof. Whether organisms also possess systems that protect genes from disruption-after TE-silencing systems fail and TEs mobilize into genes-is not well understood.

DNA transposons are TEs that typically possess a transposase-encoding gene flanked by inverted terminal repeat elements (*ITRs*)^6^. Surprisingly, insertion of DNA transposons into plant or animal genes does not always destroy host-gene function, despite the TE-based introduction of premature stop codons (PTCs) into these genes^7–10^. A likely explanation for this paradox was suggested by the observation that DNA transposons can be imprecisely excised from mRNAs in animals^7^. We speculated that the excision of TEs from mRNA was an active, host-mediated process that evolved to protect genes from TE-mediated disruption and we set out to explore this possibility.

RSD-3 is an epsin N-terminal homology (ENTH) domain protein required for RNA interference (RNAi) in *C. elegans*^11^. RNAi-mediated depletion of the *dpy-6* mRNA causes an RSD-3-dependent Dumpy (Dpy) phenotype (Fig. 1a). *Tc1* is the most active and abundant DNA transposon in *C. elegans*^12^. A *Tc1* insertion in the first coding exon of *rsd-3* ((*rsd-3*(*pk2013*), henceforth *Tc1::rsd-3*)) does not abolish *rsd-3* function in RNAi (Fig. 1a)^9^, despite introduction of PTCs, because *Tc1* is excised from *rsd-3* mRNA^9^. We first used nanopore long-read sequencing to verify that *Tc1* is excised from the *rsd-3* mRNA (Fig. 1b). We also sequenced RNA from animals in which *Tc1* has been mobilized into coding exons of five other *C. elegans* genes to assess generality ((Extended Data (ED) Fig. 1)). To capture all *Tc1* excision events, including those that created out-of-frame mRNAs, RNA from *smg-2*(−);*Tc1::rsd-3* animals, which are defective for nonsense-mediated decay^13^, was sequenced (Fig. 1b). Finally, *in vitro* synthesized *rsd-3* RNAs, which did or did not contain *Tc1*, were included to ensure that *Tc1* excision was not an *in vitro* artifact of library preparation or sequencing (Fig. 1b). The analysis showed that in all cases *Tc1* was efficiently excised from 90-100% of its host mRNAs *in vivo* (Fig. 1b and ED Fig. 1). Also in all cases, *Tc1*, which exhibits a low rate of transposition from DNA^14^, was present in DNA but not mRNA in these animals (Fig. 1c and ED Fig. 1), indicating that *Tc1* excision had occurred at the level of RNA. TE excision from mRNA exhibited the following properties, which support and extend previous studies^7–9^; 1) Excision occurs at multiple sites in or near *Tc1 ITRs* (Fig. 1d); 2) A subset of the *Tc1*-excised mRNAs are in-frame (Fig. 1d); 3) Excision sites only rarely map to consensus spliceosomal GU-AG^15^ splice sites (Fig. 1e); and 4) Excision leaves short insertions or deletions (indels) in repaired mRNAs (Fig. 1d). Henceforth, we refer to the removal of a TE from its host mRNA as SOS splicing.

**Fig. 1.**
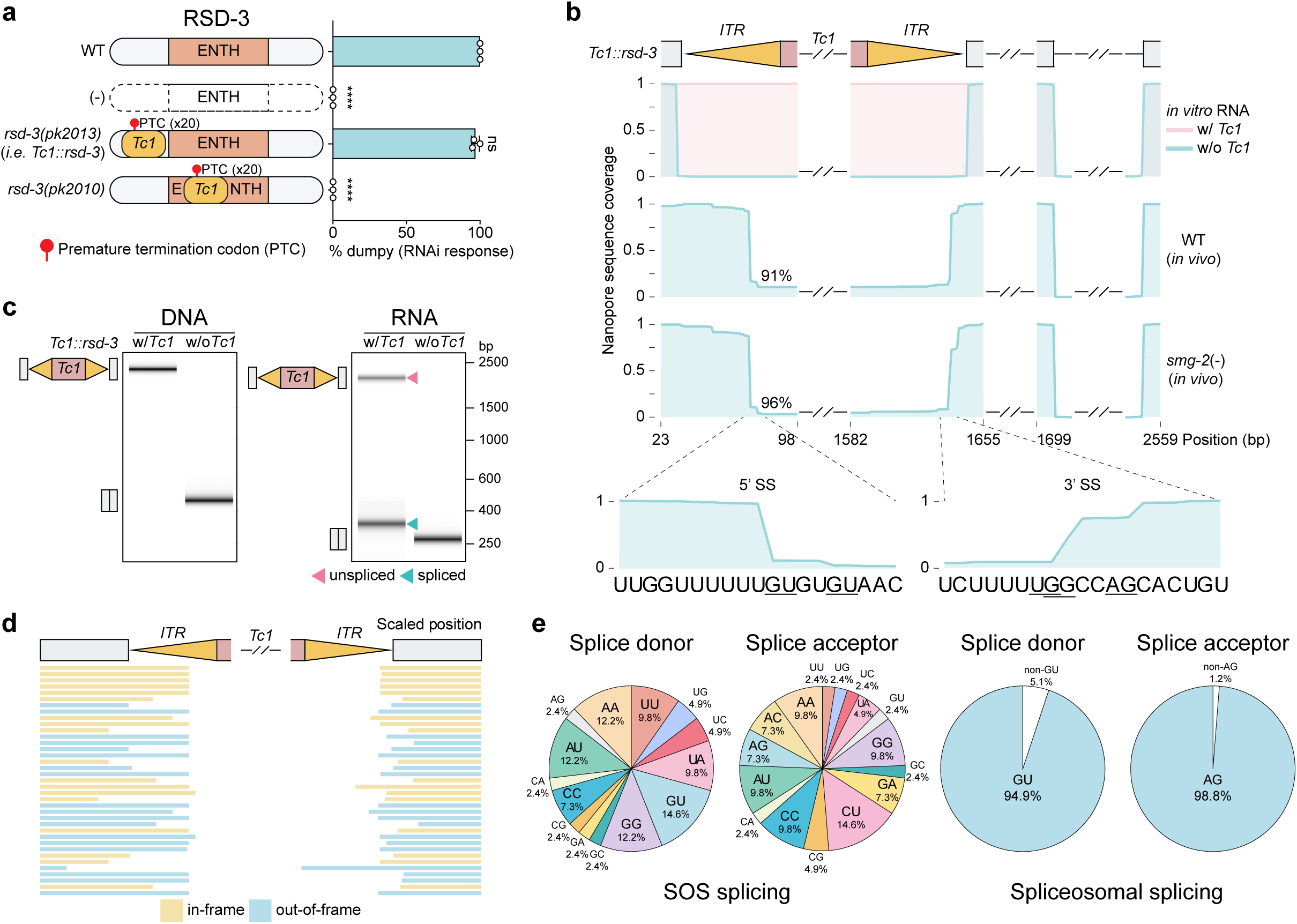
SOS splicing excises DNA transposons from *C. elegans* mRNAs. **a.** Wild-type, *rsd-3* deletion, or animals harboring the indicated *Tc1* insertions, were treated with *dpy-6* dsRNA (RNAi). The % of animals exhibiting a dumpy (Dpy) phenotype caused by *dpy-6* RNAi knockdown is shown. Data are presented as mean ± SD with all data points shown. *N* = 3 biologically independent experiments. One-way ANOVA analysis followed by Tukey’s post-hoc test. \*\*\*\**P* < 0.0001, ns not significant, *P* > 0.05. **b.** Nanopore long-read sequencing of *in vitro* synthesized *rsd-3* RNAs with or without *Tc1*, and RNA isolated from animals of indicated genotypes. PCR amplicons for sequencing were generated with primers flanking the *Tc1* insertion site in *rsd-3*. SOS splicing efficiency is indicated. Sequences bordering 5’ and 3’ SOS splice sites (SS) are shown, with common SOS splice sites underlined. *Tc1::rsd-3* is *rsd-3(pk2013*). **c.** SOS splicing visualized by automated DNA electrophoresis via Agilent Tapestation (henceforth, Tapestation). PCR amplicons were generated from genomic DNA (left panel) or cDNA (RT-PCR) (right panel). Wild-type (N2) (w/o *Tc1*) serves as a negative control. Unspliced and spliced amplicons are indicated. *Tc1::rsd-3* is *rsd-3(pk2013*). **d.** Summary of all SOS splicing events detected in *Tc1::rsd-3 ((rsd-3(pk2013))*, *smg-2*(−);*Tc1::rsd-3,* and five other *C. elegans* exonic *Tc1* insertions (see ED Fig. 1). In-frame and out-of-frame isoforms are highlighted in yellow and blue, respectively. **e.** (Left) The two nucleotides bordering SOS splice sites for *C. elegans smg-2(−);Tc1::rsd-3* (this figure), *C. elegans Tc1-traΔ::ITRscr::rsd-3* (see Fig. 2), and *HSMAR2-GFP* in human HEK293T cells (see Fig. 4) are shown. (Right) Nucleotides bordering spliceosomal splice sites for 10,000 randomly selected *C. elegans* introns.

**Fig. 2.**
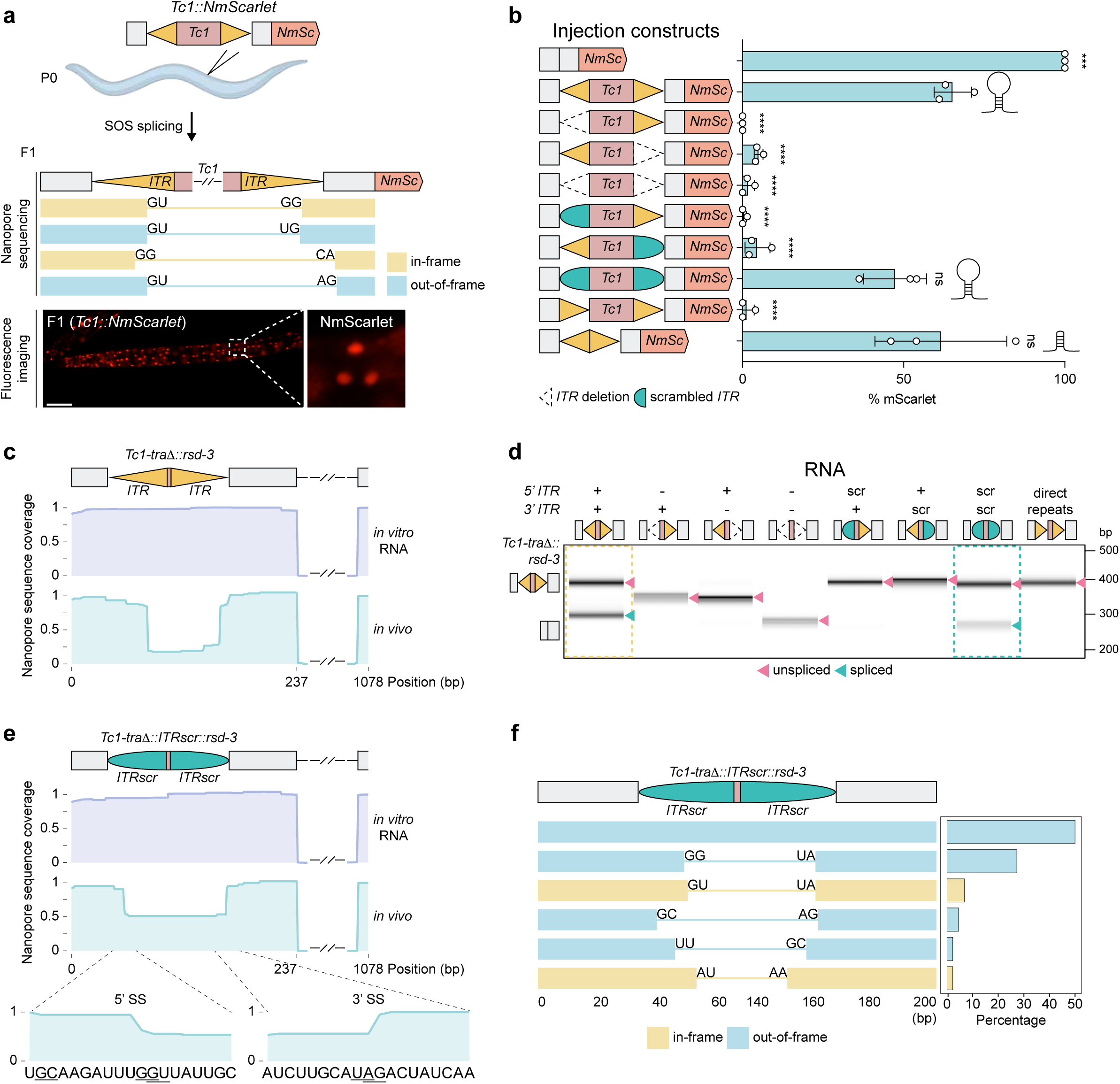
SOS splicing is a pattern recognition system triggered by RNA structure. **a.** (Top) Schematic of assay to explore SOS splicing in *C. elegans.* Plasmids harboring *Tc1::NmScarlet* (nuclear mScarlet) were injected into adult *C. elegans* (P0). (Middle) SOS splicing was visualized using nanopore sequencing of RNA isolated from F1 progeny. (Bottom) SOS splicing was associated with mScarlet expression in F1 progeny. **b.** Indicated constructs were injected and % of F1 progeny expressing mScarlet signal was quantified. (Left) Schematic of injected constructs, which all contain NmScarlet. Variants retaining the ability to engage in *ITR* base-pairing are indicated by hairpin schematics. Data are presented as mean ± SD with all data points shown. *N* = 3 biologically independent experiments. One-way ANOVA analysis followed by Tukey’s post-hoc test. \*\*\**P* < 0.001, \*\*\*\**P* < 0.0001, ns not significant, *P* > 0.05. **c.** Nanopore sequencing of RNA isolated from *Tc1-traΔ::rsd-3* animals. *In vitro* synthesized RNAs are included as controls. **d.** SOS splicing of chromosomally integrated *Tc1-traΔ* reporter variants detected with Tapestation. (Top) Schematic of variants. RT-PCR amplicons for indicated *Tc1-traΔ* variants are shown. Unspliced and spliced amplicons are indicated with arrows. Variants undergoing SOS splicing are indicated with dashed rectangles. Scr, scrambled *ITR*. **e.** Nanopore sequencing of RNA isolated from *Tc1-traΔ::ITRscr::rsd-3* animals is shown*. In vitro* synthesized *Tc1-traΔ::ITRscr::rsd-3* RNA is shown. **f.** SOS splicing isoforms observed in (**e**), ranked by abundance. Isoforms representing > 2% of total reads are shown.

Analysis of additional *Tc1* insertions revealed additional features of SOS splicing. First, *Tc1* located in the 3’ UTR of an mRNA was subjected to SOS splicing, however, two intronic *Tc1* insertions were not (ED Fig. 2). Second, in two cases in which *Tc1* was located near (<11 nt) an intron-exon boundary of its host exon, *Tc1* was removed, at least partly, from its host mRNA via apparent hybrid splicing between canonical 5’ or 3’ spliceosome splice sites from a neighboring host-gene exon and SOS splice sites near 3’ or 5’ termini of *Tc1 ITRs*, respectively (ED Fig. 3)^7^. The data hint that proximity of a TE to an intron-exon boundary could lead to hybrid spliceosome-SOS splicing. The mechanism and provenance of hybrid spliceosome-SOS splicing is not investigated further here. Finally, when *Tc1* was inserted into eight different coding sites in the *rsd-3* gene, all insertions were subjected to SOS splicing, however, SOS splicing only rescued *rsd-3* function in RNAi when *Tc1* was inserted into regions of *rsd-3* that are poorly conserved (*i.e.* not the ENTH domain) (Fig. 1a and ED Fig. 4). Together, the data suggest that SOS splicing will operate on any *Tc1* element located in mRNA, but that SOS splicing will only rescue gene function when the indels left in repaired mRNAs by SOS splicing do not disrupt essential protein functions.

We wondered how *C. elegans* might identify *Tc1*-containing RNAs to initiate SOS splicing. We engineered a mScarlet gene containing a *Tc1* element (*Tc1::NmScarlet*), which, if subjected to SOS splicing, would produce nuclear-localized mScarlet (Fig. 2a). Injection of this reporter gene into the germline of adult *C. elegans* resulted in progeny that expressed mScarlet, suggesting SOS splicing had occurred (Fig. 2a). RNA sequencing confirmed that the *Tc1::NmScarlet* mRNA was SOS spliced (Fig. 2a). Deleting all sequences, including transposase, located between the *ITRs* of *Tc1::NmScarlet* did not interfere with *Tc1::NmScarlet* expression, indicating dispensability of these sequences for triggering SOS splicing (Fig. 2b). Similarly, deletion of the transposase gene from a genomic *Tc1::rsd-3* locus (*Tc1-traΔ::rsd-3*) did not impact SOS splicing, as indicated by nanopore sequencing (Fig. 2c), RT-PCR analysis (Fig. 2d and ED Fig. 5), and RSD-3 functional analyses (ED Fig. 5). However, deleting either or both 54 nucleotide *ITR* (*ITRΔ*) from *Tc1::NmScarlet* abrogated mScarlet expression (Fig. 2b), and deleting an *ITR* element from chromosomally integrated *Tc1::rsd-3, Tc1-traΔ::rsd-3*, or *Tc1::unc-54* genes prevented SOS splicing of *Tc1* from its host mRNAs (Fig. 2d and ED Fig. 5). We conclude that *ITRs* are necessary for SOS splicing.

*ITRs* could base-pair to form dsRNA hairpin structures, which might be the signal that triggers SOS splicing. Consistent with this idea, SOS splicing of *Tc1::NmScarlet* required that *ITRs* be inverted with respect to each other (Fig. 2b). To test this idea further, we scrambled the *Tc1 ITR* sequence (*ITRscr*) and replaced the *ITR* elements of *Tc1::NmScarlet* or *Tc1-tra*Δ*:rsd-3* with *ITRscr* in an inverted orientation. Thus, in these *ITRscr* genes, the *ITR* sequence is scrambled, however, scrambled *ITRs* retain the ability to base-pair with each other. mScarlet expression was observed in *Tc1-ITRscr::NmScarlet* animals (Fig. 2b), and nanopore sequencing (Fig. 2e-f) and RT-PCR analysis (Fig. 2d) revealed that SOS splicing occurred on the *Tc1-tra*Δ*::ITRscr*::*rsd-3* mRNA, albeit with reduced efficacy when compared to wild-type *Tc1*. The data suggest that SOS splicing is a pattern recognition system triggered by the base-pairing of inverted repeats, which are a defining feature of the DNA transposons.

If SOS splicing is an active host response to a TE interrupting an mRNA then a genetic screen could identify host factors mediating SOS splicing. We generated *C. elegans* expressing two SOS splicing reporter genes; *Tc1::Ngfp* and *Tc1::rsd-3*. *Tc1::Ngfp*;*Tc1::rsd-3* animals expressed GFP and were competent for RNA interference (RNAi) (see below), because both TE-containing mRNAs were subjected to SOS splicing (see Fig. 1b, and ED Fig. 3). *utp-20* RNAi causes larval arrest^16^. We mutagenized *Tc1::Ngfp*;*Tc1::rsd-3* animals and screened ≅100,000 haploid genomes to identify twenty mutant animals that failed to arrest development when subjected to *utp-20* RNAi and that did not express GFP (Fig. 3a). Genome sequencing identified candidate SOS splicing genes in the twenty mutants: We identified eight mutations in the gene *c07h6.8*, eleven mutations in *sut-2*, and one mutation in *f15d4.2* (Fig. 3b). The chromosomal locations of *c07h6.8*, *sut-2*, and *f15d4.2* were consistent with positional mapping data (ED Fig. 6). CRSPR-Cas9-based introduction of stop codons into *c07h6.8*, *sut-2,* or *f15d4.2* resulted in animals that failed to express GFP or RSD-3 from *Tc1::Ngfp* or *Tc1::rsd-3*, respectively (Fig. 3c). These mutants were also defective for SOS splicing of *ITR*-containing mRNAs (Fig. 3d and ED Fig. 7), confirming that C07H6.8, SUT-2, and F15D4.2 are required for rescuing the function of TE-interrupted genes via SOS splicing. We refer to *c07h6.8* and *f15d4.2* henceforth as *akap-17* and *caap-1*, respectively, for reasons outlined below. *akap-17* encodes a protein with an RS domain, two protein kinase A-anchoring domains (PBD), and an RNA recognition motif (RRM) (Fig. 3b). The putative mammalian ortholog of AKAP-17 is <u>A</u>-<u>k</u>inase <u>a</u>nchoring <u>p</u>rotein <u>17A</u> (AKAP17A/XE7), which is a nuclear speckle-localized protein linked to alternative splicing of reporter minigenes in human cells^17,18^. The putative mammalian ortholog of CAAP-1 is CAAP1, which promotes chemotherapeutic resistance^19–21^ and may^21,22^ or may not^23^ regulate apoptosis. The putative mammalian ortholog of SUT-2 is mSUT2/ZC3H14/NAB2, which is a conserved polyA RNA-binding and -regulating protein^24,25^ linked to tauopathy resistance^26,27^ and circular RNA biogenesis^28^. The role of SUT-2 in SOS splicing is not explored further in this work. We conclude that AKAP-17, SUT-2, and CAAP-1 are conserved proteins required for SOS splicing in *C. elegans*.

**Fig. 3.**
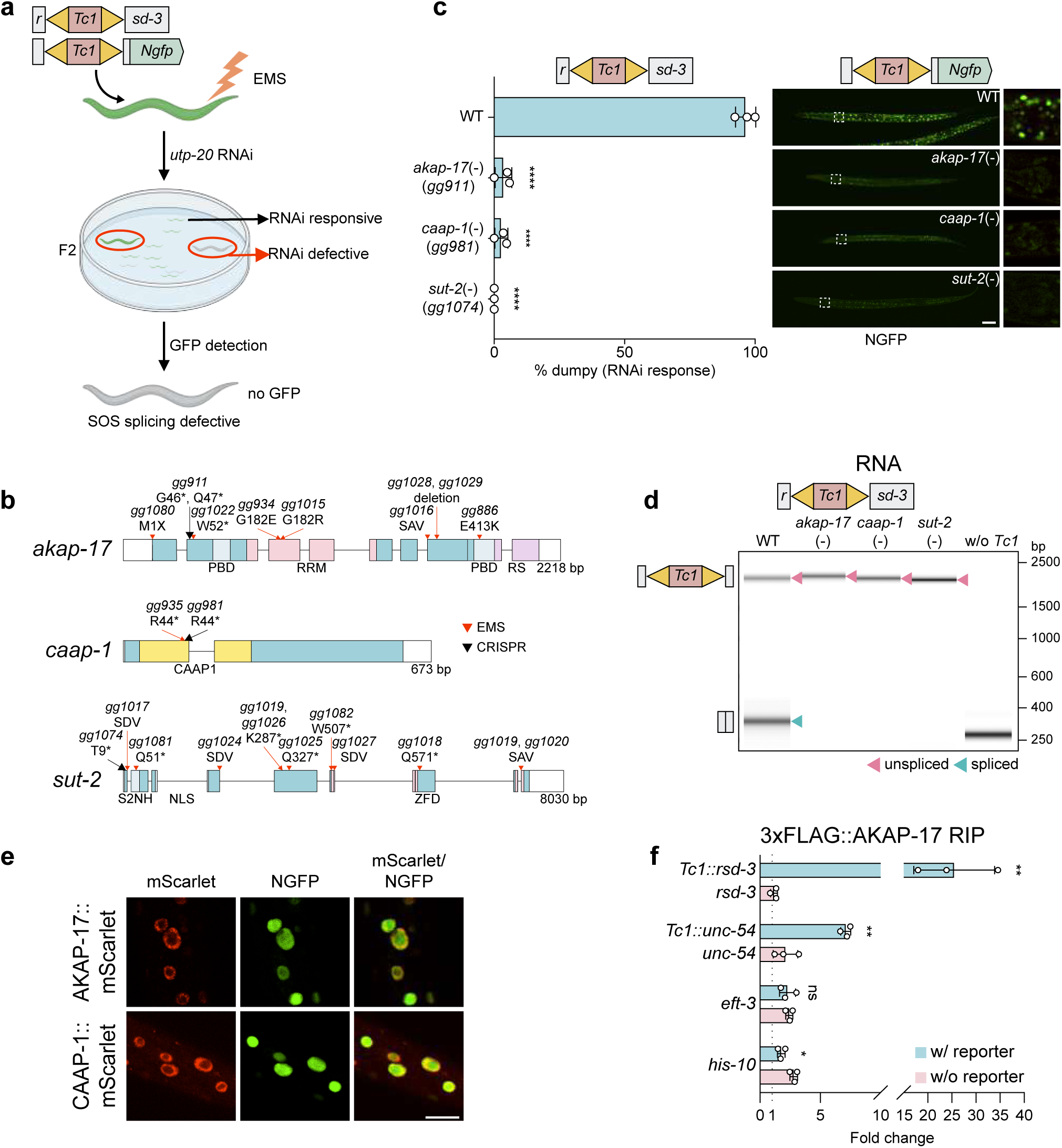
Identification of three *C. elegans* factors required for SOS splicing. **a.** Schematic of genetic screen used to identify SOS splicing factors. Animals harboring *Tc1::rsd-3*; *Tc1::Ngfp* (nuclear GFP) were mutagenized with ethyl methanesulfonate (EMS). F2 progeny from mutagenized animals were treated with *utp-20* dsRNA. Mutant animals that failed to arrest development in response to *utp-20* RNAi were isolated. Lineages established from these animals were then screened for lack of GFP expression. **b.** Alleles of *akap-17*, *caap-1* and *sut-2* identified in screen are indicated with red arrows. Alleles generated by CRISPR-Cas9 are indicated with black arrows. Predicted protein domains are shown. PBD: protein kinase A-anchoring domain; RRM: RNA recognition motif; RS: arginine/serine-rich domain; CAAP1: caspase activity and apoptosis inhibitor 1; S2NH: SUT-2 N-terminal homology; NLS: nuclear localization signal; ZFD: zinc finger domain. *: stop codon; SDV: splice donor variant; SAV: splice acceptor variant; M1X: initiating methionine mutation. **c.** (Left) RNAi responsiveness and (Right) GFP expression in animals of the indicated genotypes, and harboring the indicated reporter genes, is shown. (Right) Boxed regions are magnified and shown to the far right. Scale bar = 50 μm. Data are mean ± SD with all data points shown. (−) indicates nonsense alleles, which are indicated in left panels, were assayed. *N* = 3 biologically independent experiments. One-way ANOVA analysis followed by Tukey’s post-hoc test. \*\*\*\**P* < 0.0001. **d.** Tapestation-based demonstration that candidate SOS splicing factors are defective for SOS splicing. (−) indicates *akap-17(gg911)*, *caap-1(gg981)*, and *sut-2(gg1074)*, respectively. Unspliced and spliced amplicons are indicated. All animals express *Tc1::rsd-3*. **e.** Spinning disc confocal images of somatic cells from larval stage four *C.elegans* in which mScarlet was introduced into *akap-17* or *caap-1* genes (see ED Fig. 8 for details). These animals also express the *Tc1::Ngfp* SOS reporter gene, which produces Nuclear GFP (NGFP). The data show that; 1) AKAP-17/CAAP-1 are nuclear proteins; and 2) that SOS splicing is functional in *akap-17* and *caap-1* tagged animals. Scale bar = 10 μm. **f.** AKAP-17 RNA immunoprecipitation (RIP) analysis. FLAG::AKAP-17 was immunoprecipitated using anti-FLAG. AKAP-17-bound RNAs were quantified by qRT-PCR. Data was normalized to RIP signals from animals expressing AKAP-17 without FLAG tag. Data are presented as mean ± SD. All data obtained, after optimizing RIP protocol, are shown. *N* = 3 biologically independent experiments. Two-tailed unpaired student’s *t*-test. \**P* < 0.05, \*\**P* < 0.01, ns not significant, *P* > 0.05.

We introduced N-terminal mScarlet tags to *C. elegans akap-17* and *caap-1*. The tags did not impact AKAP-17 or CAAP-1 function as these animals were proficient for SOS splicing (ED Fig. 8). mScarlet::AKAP-17 and mScarlet::CAAP-1 fluorescence was observed in nuclei of all/most *C. elegans* cells in both the soma and germline and at all stages of development (Fig. 3e, and ED Fig. 8). The data suggest that SOS splicing is a nuclear process that could be active in all *C. elegans* cell types.

mRNA sequencing of rRNA-depleted RNA from wild-type or *akap-17*(−) animals showed that most spliceosome-mediated splicing events occurred normally in *akap-17*(−) animals (ED Fig. 8). Thus, the role of AKAP-17 in RNA splicing is relatively specific to SOS splicing. Given this specificity, and because AKAP-17 has an RRM domain, we asked if AKAP-17 might interact with TE-containing mRNAs. Indeed, AKAP-17 RNA IP (RIP) analysis showed that AKAP-17 co-precipitated with two mRNAs whose parent genes contained *Tc1*, but not four control mRNAs lacking *Tc1* (Fig. 3f). The association of AKAP-17 with TE-containing mRNAs suggests AKAP-17 could participate directly in the SOS splicing process.

Given that AKAP-17, SUT-2, and CAAP-1 are conserved in mammals, we wondered if SOS splicing might also occur in human cells. To address this question, we constructed plasmids that express a GFP gene interrupted by either *C. elegans Tc1* or the human *hsMar2* DNA transposon (*Tc1-GFP* or *HSMAR2-GFP*). These reporter genes would not be expected to produce GFP, due to introduced PTCs, unless an SOS splicing system was operational in human cells. Transfection of either plasmid into human embryonic kidney (HEK)-293T cells resulted in cells that expressed GFP (Fig. 4a). RT-PCR analysis (ED Fig. 9) and nanopore sequencing (Fig. 4b-c) showed that both transposons were excised from their host mRNAs in human cells. The molecular hallmarks of TE excision in human cells resembled those of SOS splicing in *C. elegans*; TE excision was efficient and was not restricted to spliceosome associated GU-AG splice sites (Fig. 4c and ED Fig. 9). Interestingly, while excision sites for both *Tc1* and *HSMAR2* occurred near their respective *ITRs*, the precise sites of excision differed; *Tc1* excision occurred within *Tc1 ITRs* and *HSMAR2* excision sites occurred 100-393 nt distal to *HSMAR2 ITRs* (Fig. 4b and see discussion). The data suggest that a SOS splicing-like system is active in human cells.

**Fig. 4.**
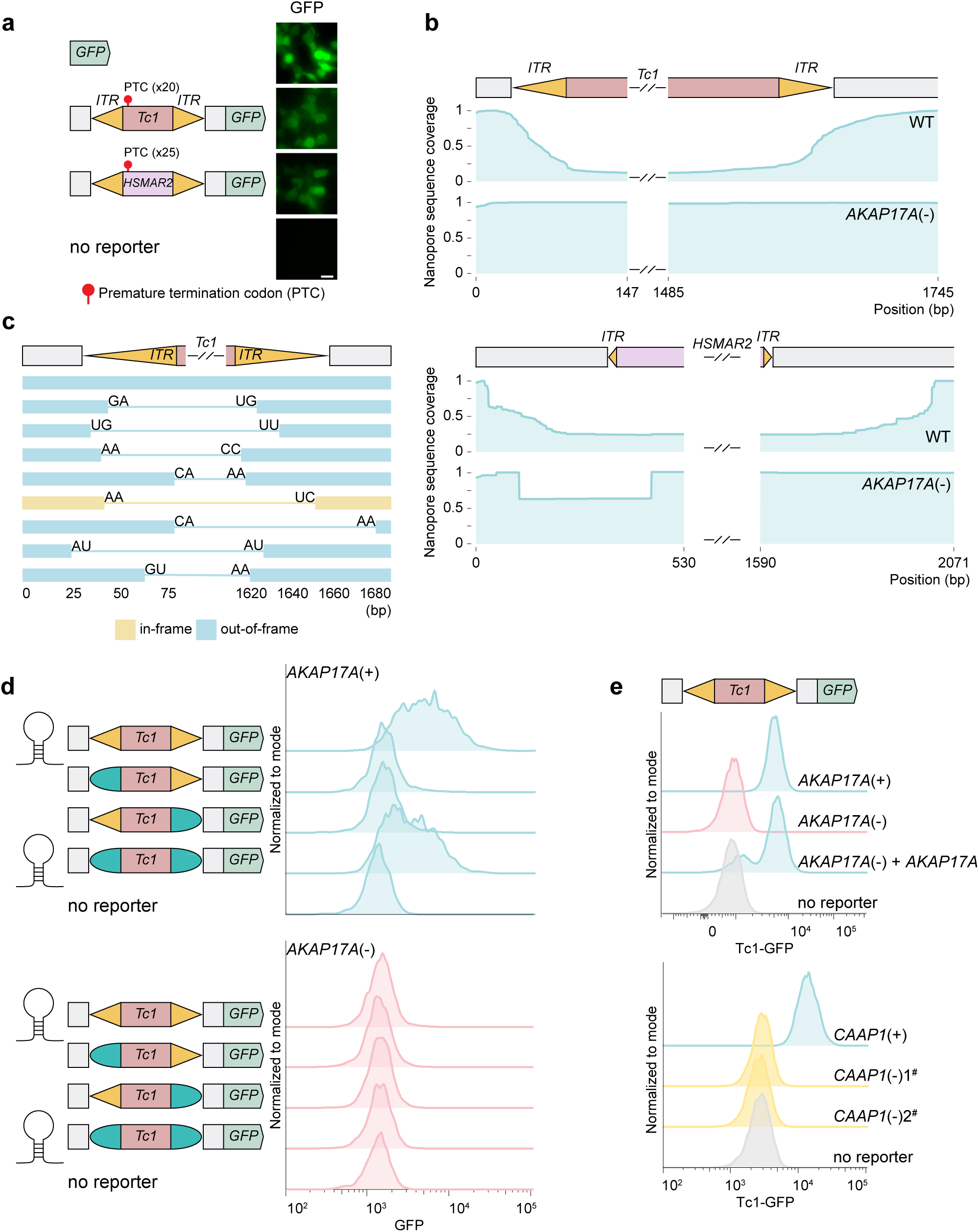
SOS splicing in human cells. **a.** Representative fluorescence micrographs of HEK293T cells transfected with indicated human SOS splicing reporter plasmids. (Left) Schematic of transfected reporter constructs. Scale bar = 20 μm. **b.** Nanopore sequencing of RNA isolated from *AKAP17A*(+) (WT) *or* (−) HEK293T cells transfected with (Top) *Tc1-GFP* SOS splicing reporter or (Bottom) *HSMAR2-GFP* reporter. See ED Fig. 9 for data demonstrating cells are *AKAP17A*(−). (Bottom) The source of the small deletion seen in *HSMAR2-GFP* RNAs in *AKAP17A*(−) cells is not known. **c.** SOS splicing isoforms detected in *Tc1-GFP* mRNA from WT HEK293T cells, ranked by abundance. Isoforms representing > 0.5% of total reads are shown. **d.** Flow cytometry of *AKAP17A*(+) and *AKAP17A*(−) HEK293T cells transfected with the indicated *Tc1-GFP* variant constructs. Non-transfected cells serve as a negative control. *Tc1-GFP* constructs that retain *ITR* base-pairing capability are indicated. (Left) Schematic of variant reporters. **e.** Flow cytometry of HEK293T cells harboring a chromosomally integrated *Tc1-GFP* SOS splicing “knock-in” reporter gene. (Top) *AKAP17A*(+) or (−) cells, and *AKAP17A*(−) cells complemented with *AKAP17A(+)* via lentiviral transformation, are shown. (Bottom) *CAAP1*(+) cells, and two independently derived clones (1# and 2#) of *CAAP1*(−) HEK293T cells, are shown. See ED Fig. 9 for data showing cells are *CAAP1*(−).

To explore this idea further, we asked if *ITR* elements were necessary and sufficient for TE excision from reporter mRNAs in human cells, as they were in *C. elegans.* Indeed, structure-function analyses of the *Tc1-GFP* reporter gene showed that; 1) the *ITRs* of *Tc1-GFP* were necessary and sufficient to elicit TE excision from *Tc1-GFP;* and 2) excision occurred independently of the underlying *ITR* sequence (Fig. 4d). We next asked if the putative mammalian orthologs of AKAP-17 or CAAP-1 were needed for TE excision in human cells. We inserted *Tc1-GFP* or *HSMAR2-GFP* reporter genes into the AAVS1 safe-harbor site^29^ in HEK293T cells. We confirmed that *Tc1-GFP* and *HSMAR2-GFP* expressed GFP using flow cytometry (Fig. 4e, ED Fig. 9). We then deleted all copies of the *AKAP17A* or CAAP1 genes from *Tc1-GFP* or *HSMAR2-GFP* cells (ED Fig. 9) and observed that *AKAP17A*(−) or *CAAP1*(−) cells no longer expressed GFP (Fig. 4e, ED Fig. 9). Reintroduction of a wild-type copy of *AKAP17A* into *AKAP17A*(−) cells, using lentiviral transduction, rescued GFP expression from *Tc1-GFP* and *HSMAR2-GFP* (Fig. 4e, ED Fig. 9). Together, the data show that SOS splicing occurs in human cells and that orthologous proteins and related RNA structures mediate SOS splicing in humans and nematodes.

The diversity of SOS splice sites, and the lack of spliceosomal GU-AG signatures at these sites, suggests that SOS splicing is not mediated by the spliceosome but, rather, by a different RNA cleavage and ligation mechanism. Because neither AKAP17A, CAAP1, or mSUT2 exhibit homology to known endonucleases or ligases, we conducted IP-mass spectrometry (MS) on human CAAP1 in an attempt to further understand the mechanism of SOS splicing. CAAP1 IP-MS from HEK293T cells identified AKAP17A and mSUT2, as well as the RNA ligase RTCB, as candidate CAAP1-interacting proteins (Fig. 5a). Some tRNAs possess introns, which are excised by the endonucleases TSEN2/4^30^. The resultant tRNA fragments are ligated by RTCB to generate mature tRNAs^31^. RTCB also ligates *XBP1* mRNA fragments generated during the unfolded protein response^32^. We wondered if RTCB might contribute to SOS splicing, perhaps by ligating mRNA fragments created by the excision of TEs during SOS splicing. Directed Co-IP analyses confirmed the CAAP1 IP-MS results: CAAP1-RTCB and CAAP1-AKAP17A co-IPed in both *C. elegans* and in human cells (Fig. 5b/c and ED Fig. 10). AlphaFold3 simulations hint that CAAP1-RTCB and CAAP1-AKAP17A may interact directly (ED Fig. 10, ipTM values 0.23-0.47). We used the tripartite split-GFP system to assess where in the cell RTCB might interact with the other SOS splicing factors^33^. We co-expressed; 1) AKAP17A fused to the 10th beta strand of GFP; 2) RTCB fused to the 11th beta strand of GFP; and 3) beta strands 1-9 of GFP in HEK293T cells. In these cells, we observed GFP fluorescence in nuclear foci, suggesting AKAP17A-RTCB interact in these foci (Fig. 5d). GFP fluorescence was not observed when GFP beta strands were not fused to AKAP17A or RTCB (Fig. 5d). Because CAAP1 associates with both RTCB and AKAP17A, we wondered if CAAP1 might be necessary for RTCB and AKAP17A to interact. Indeed, in *CAAP1*(−) cells, RTCB and AKAP17A failed to interact via split-GFP (Fig. 5d). Additionally, transiently transfected AKAP17A and RTCB could be co-IPed from HEK293T cells, but only when CAAP1 was also co-expressed in these cells (Fig. 5e). Thus, AKAP17A and RTCB interact in the nucleus and CAAP1 promotes this interaction. And because AKAP17A localizes to nuclear speckles^17^, the interaction between RTCB and AKAP17A is likely to occur in nuclear speckles. Related split-GFP analyses showed that CAAP1-AKAP17A also interact in nuclear foci and that RTCB-CAAP1 interact throughout cells (Fig. 5d). Taken together, the data suggest that SOS splicing is likely to occur in nuclear speckles in human cells and that CAAP1 recruits the RNA ligase RTCB to these foci.

**Fig. 5.**
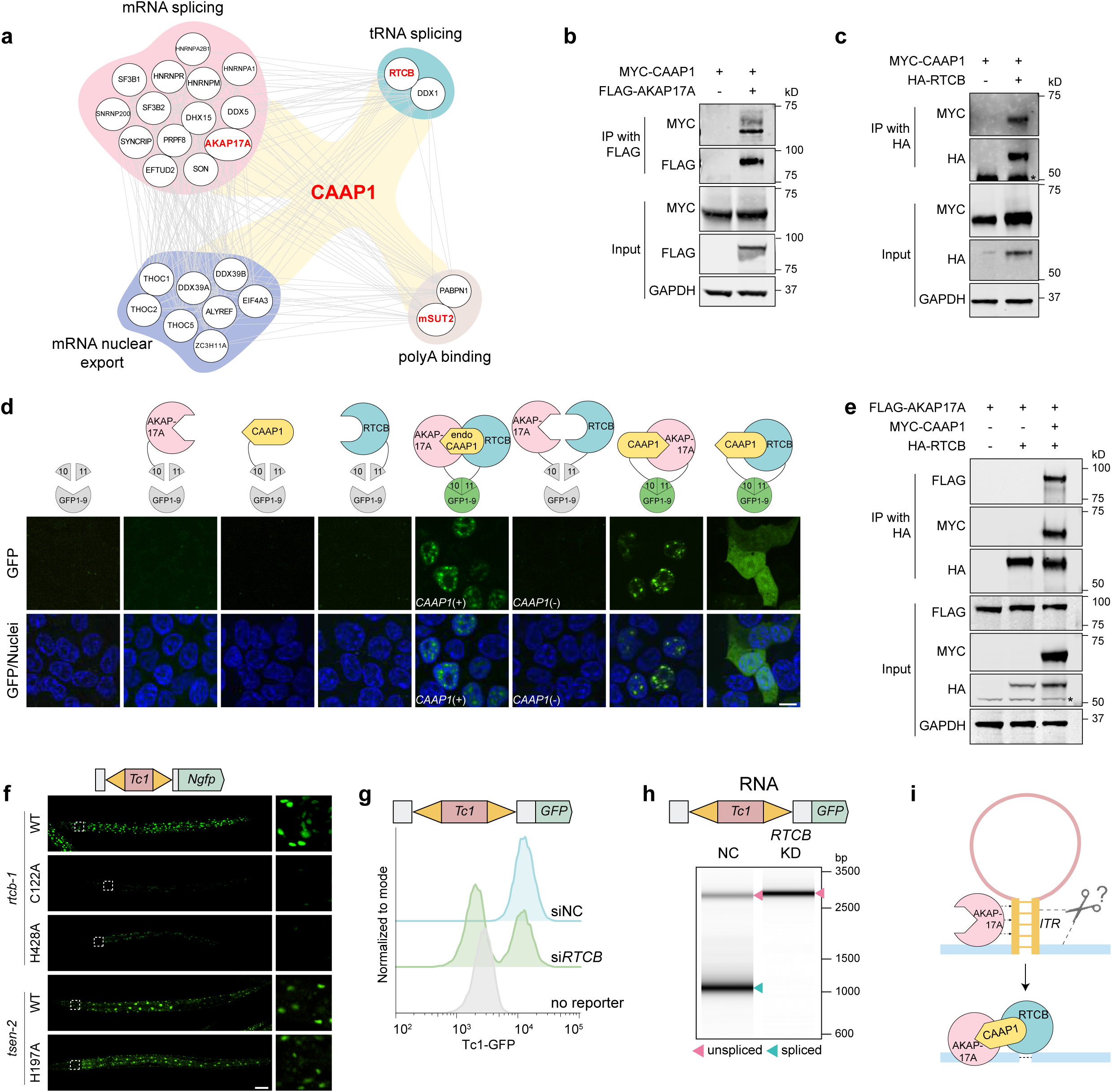
The RNA ligase RTCB is required for SOS splicing. **a.** STRING network analysis of CAAP1 Immunoprecipitation-Mass Spectrometry (IP-MS) results from HEK293T cells. Factors identified in the genetic screen, as well as the RNA ligase RTCB, are shown in red font. **b-c**. Co-immunoprecipitation (Co-IP) assay showing interactions between AKAP17A and CAAP1 (**b**), RTCB and CAAP1 (**c**) in HEK293T cells transfected with plasmids expressing the indicated tagged-proteins. GAPDH serves as a loading control. * Non-specific band. **d.** (Top) Schematic of tripartite split-GFP system. (Bottom) Representative images showing interactions between AKAP17A, CAAP1 and RTCB. Nuclei were stained with Hoechst 33342. Scale bar = 10 μm. **e.** Co-IP assay showing interactions between AKAP17A and RTCB. The AKAP17A-RTCB interaction is only observed when CAAP1 is co-transfected. GAPDH serves as a loading control. * Non-specific band. **f.** Fluorescence micrographs showing *Tc1::Ngfp* SOS splicing reporter expression in WT, RTCB-1(C122A), RTCB-1(H428A), or TSEN-2(H197A) animals. Boxed regions are magnified to the right. Scale bar = 50 μm. **g.** Flow cytometry of GFP expression in HEK293T *Tc1-GFP* reporter knock-in HEK293T cells treated with esiRNA targeting *RTCB* (si*RTCB*) or Renilla Luciferase (siNC). **h.** Tapestation analysis of amplicons generated from RNA extracted from HEK293T cells treated with esiRNA targeting RTCB (KD) or Renilla Luciferase (NC) and transfected with *Tc1-GFP* reporter plasmid. Unspliced and spliced amplicons are indicated with arrows. **i.** Model for SOS splicing. Scissors represent an unknown endonuclease that removes TEs from mRNAs. It is not yet clear if AKAP17A interacts directly with *ITR* hairpins, as indicated by the model.

Finally, we asked if RTCB was required for SOS splicing. Note that RTCB is essential for viability, due to its role in tRNA splicing, and, therefore, would not likely have been identified by our genetic screen. We used CRISPR-Cas9 to alter two residues in *C. elegans* RTCB ((RTCB-1(C122A, H428A)), which are required for RTCB-based RNA ligation^34,35^. Heterozygous C122A/+ or H428A/+ animals were isolated and their C122A or H428A homozygous progeny were found to arrest at larval stage three (L3) of development (ED Fig. 10). Homozygous C122A or H428A progeny, which harbored the *Tc1::Ngfp* SOS splicing reporter gene, failed to express GFP as L2/L3 animals, suggesting RTCB-1-based RNA ligation is required for SOS splicing (Fig. 5f). Similar results were obtained with animals that expressed the *Tc1::NmScarlet* SOS splicing reporter gene and that were homozygous for a deletion allele of *rtcb-1* (ED Fig. 10). Animals harboring a catalytic site mutation in TSEN-2 (tRNA intron endonuclease) arrested development similarly to *rtcb-1* mutants (L4 stage) (ED Fig. 10), however, these animals did not exhibit defects in SOS splicing (Fig. 5f), suggesting that; 1) the SOS splicing defects observed in *rtcb-1* mutants are not an indirect consequence of failure to splice tRNAs; and 2) that the TE excision step of SOS splicing is mediated by another, currently unknown endonuclease. Finally, siRNA-based knockdown of *RTCB* in HEK293T cells decreased expression of GFP from *Tc1-GFP* or *HSMAR2-GFP* (Fig. 5g and ED Fig. 10) and RT-PCR analysis showed that SOS splicing of *Tc1-GFP* and *HSMAR2-GFP* became inefficient when RTCB was depleted (Fig. 5h and ED Fig. 10). The data show that the RNA ligase RTCB promotes SOS splicing in *C. elegans* and human cells. The data suggest that in human cells RTCB is recruited to nuclear speckles by CAAP1 where it interacts with AKAP17A to promote SOS splicing by ligating mRNA fragments generated during the TE-excision step of SOS splicing.

## Discussion

Here we describe a new mode of mRNA splicing, which we term SOS splicing. We show that SOS splicing is a conserved pattern recognition system that detects inverted repeats in mRNAs and excises them. We identify four proteins required for SOS splicing; the TE::mRNA binding AKAP17A; the RNA ligase RTCB; CAAP1, which recruits RTCB to AKAP17A in nuclear foci; and mSUT2, whose role in SOS splicing has not yet been investigated (Fig. 5i). Finally, we show that one biological function of SOS splicing is to remove DNA transposons from mRNAs, thus restoring function to some TE-interrupted genes.

We propose that RTCB is the enzyme that ligates mRNA fragments generated during SOS splicing because RTCB-based RNA ligation is required for SOS splicing and because RTCB interacts physically with the other SOS splicing factors. We do not yet understand how TEs are excised from mRNAs, prior to RTCB ligation. The tRNA intron endonuclease TSEN-2 does not appear to be required for SOS splicing, suggesting a different cleavage mechanism is at play. While SOS splicing can be >99% efficient, it is not precise. For example, while SOS splice sites occur near the 3’ or 5’ termini of *ITRs*, the exact site of splicing can differ: in human cells, *Tc1* excision occurs within *ITR* elements while *HSMAR2* excision occurs ≅100-400 nt distal to its *ITRs*. Thus, although SOS splicing appears to operate on any exonic *ITR*, sequence elements found in or around *ITRs* may help dictate precisely where TE excision will occur. The imprecision of SOS splicing suggests that the enzyme- or ribozyme-that excises TEs from mRNAs is not sequence specific. AKAP17A, CAAP1, or MSUT2 may help direct RNA cleavage by this unknown nuclease.

SOS splicing did not occur when *ITRs* were located in introns, suggesting that SOS splicing is either aware of exon-intron boundaries or that spliceosomal splicing is more efficient than SOS splicing and, therefore, removes TE-containing introns before they can be subjected to SOS splicing. Only a subset of SOS spliced RNAs are in frame and SOS spliced mRNAs possess indels, indicating that SOS splicing will not fully mask the negative impacts of TE insertions for many genes. Indeed, coding TE insertions are rare in the laboratory wild-type strain of *C. elegans*, despite SOS splicing, suggesting that most coding TE insertions are sub-optimal and are purged by natural selection. On the other hand, other wild isolates of the genus *Caenorhabditis* do possess coding TE insertions, and these animals do not exhibit phenotypes associated with inactivation of TE-interrupted genes^36^, hinting that SOS splicing protects these genes, and that, in some cases, SOS splicing can enable co-existence with genic TEs insertions over evolutionary time-scales. Indeed, it has been suggested that some modern-day introns could be evolutionary descendants of ancient genic transposon insertions^37^. It is perhaps interesting to speculate that, by efficiently rescuing the function of some TE-interrupted genes, SOS splicing may have facilitated the evolution of some ancient DNA transposons into introns.

We have not yet comprehensively identified all substrates of SOS splicing. While DNA transposons are not currently active in the human genome^38^, relics of older DNA transposons constitute 2-3% of the genome^39^. It will be of interest to ask if any of these TE relics are present in human mRNAs and if they are excised by SOS splicing. In addition, all genomes contain repetitive sequences, many of which are inverted and transcribed and could, therefore, be targets of SOS splicing. Similarly, some RNA and DNA viruses possess inverted repeats, which could be recognized and processed by SOS splicing^40,41^. Because AKAP-17 associates with RNAs destined for SOS splicing, identifying the RNAs bound by AKAP17A could identify additional endogenous or exogenous targets of SOS splicing, should they exist. Determining *ITR* length and complementarity requirements for triggering SOS splicing, and assessing local sequence contributions to SOS splicing, could also help identify SOS splicing targets and perhaps suggest strategies to artificially trigger SOS splicing in human mRNAs for therapeutic benefit.

The RNA ligase RTCB is present in all three kingdoms of life^35,42,43^. DNA transposons are also present in the three kingdoms, suggesting that TEs were an ancient threat to life on earth and hinting that the original function of RTCB may have been in SOS splicing and TE mitigation. Assessing if archaea and prokaryotes possess RTCB-dependent SOS splicing systems will be a strong test of this idea. Organisms possess sophisticated systems to prevent TE expression and replication^1–4^. This study, along with other recent studies^44,45^, show that organisms also possess fail-safe systems that enable coexistence with TEs when TE-silencing systems fail and TEs mobilize into genes. Given the diversity and near-ubiquity of TEs in genomes and across evolutionary time, we speculate that additional TE-coexistence systems likely exist and await discovery.

## Methods

### C. elegans husbandry

All *Caenorhabditis elegans* (*C. elegans*) strains (Supplementary Table 1) were maintained at 20 °C on Nematode Growth Medium (NGM) plates seeded with *Escherichia coli* (*E. coli*) OP50, according to previously described methods^1^, unless otherwise specified. Strains obtained from the *Caenorhabditis* Genetics Center are listed in Supplementary Table 1. WT or wild-type refers to the N2 Bristol strain. All strains are available upon reasonable request.

### Cell lines

HEK293T cells (ATCC, CRL-3216) and HEK293T-derived reporter and knockout cell lines (Supplementary Table 1) were cultured in Dulbecco’s modified Eagle’s medium (DMEM) (Gibco, 11995-065) supplemented with 10% fetal bovine serum (FBS) (Gibco, 10437-028). Cells were maintained at 37 °C in a humidified atmosphere with 5% CO2.

### RNAi

#### Feeding RNAi

Gravid adults were treated with Alkaline-bleach solution (20% commercial bleach, 0.5 M NaOH) to isolate embryos (egg preparation). The embryos were then transferred to RNAi plates (NGM plates containing 1 mM IPTG and 25 mg/mL carbenicillin) seeded with *E. coli* HT115 expressing either control dsRNA (L4440 empty vector) or dsRNAs targeting specific genes (*utp-20*, or *dpy-6*). The *utp-20*, and *dpy-6* RNAi colonies are from the Ahringer library and all plasmid identities were confirmed by Sanger sequencing.

#### esiRNA treatment

Dried esiRNA oligos targeting *RTCB* (Sigma Aldrich, EHU009301-20UG) were resuspended in TE buffer (10 mM Tris-HCl pH 8.0, 1 mM EDTA). esiRNA targeting *RLUC* (Sigma Aldrich, EHURLUC-50UG) was used as a negative control in all RNAi experiments. HEK293T cells were transfected with esiRNAs using Lipofectamine™ RNAiMAX Transfection Reagent (Invitrogen, 13778075), following manufacturer’s protocols. Transfections were performed at a final concentration of 10 pmol. Knockdown efficiency of target genes was confirmed by immunoblot analysis.

### *In vitro* RNA synthesis

To prepare *in vitro* RNAs for controls for Nanopore long-read sequencing experiments, RNAs were synthesized using MEGAscript T7 Transcription Kit (Invitrogen, AM1334). DNA fragments were cloned into a pDONR221 plasmid (*Tc1::rsd-3*, *rsd-3*, *Tc1::Ngfp*) containing the T7 promoter using Gateway^®^ BP Clonase^TM^ II Enzyme Mix (Invitrogen, 11789-100) or pUC57 plasmid (*Tc1-traΔ::rsd-3*, *Tc1-traΔ::ITRscr::rsd-3*) using AccI (NEB, R0161S) and AvaI (NEB, R0152S), linearized, and purified using phenol-chloroform extraction to serve as DNA templates for *in vitro* transcription. Transcription reactions were incubated overnight at 37 °C. Resulting RNA was purified using the RNA Clean & Concentrator-5 (Zymo Research, R1014) and stored at −80 °C.

### Long-read nanopore sequencing and SOS splicing analysis

Total RNAs were extracted from animals using RNA Clean & Concentrator-5 (Zymo Research, R1014), followed by DNase treatment to deplete genomic DNA contamination. For cDNA synthesis, 1 μg of *in vivo* RNA (or 1 ng of *in vitro* RNA) was reverse transcribed using Induro^®^ Reverse Transcriptase (NEB, M0681L) with Random Primer Mix (60 μM) (NEB, N0447). Briefly, RNA was mixed with Random Primer Mix and dNTPs, denatured at 65 °C for 5 minutes, and then immediately chilled on ice. Induro RT Reaction Buffer, RNase Inhibitor, and Induro Reverse Transcriptase were added to the mixture. The reaction was incubated in a thermocycler with the following conditions: 2 minutes at 25 °C, followed by 30 minutes at 60 °C, and a final hold at 95 °C for 2 minutes.

SOS splicing isoforms were amplified using primers flanking transposon insertion sites. 200 ng cDNA (from *in vivo* RNA) or 10^−5^ pg cDNA (from *in vitro* RNA) was used for PCR amplification. Reactions were performed in a 200 μL reaction volume with 20-23 cycles using Q5^®^ High-Fidelity DNA Polymerase (NEB, M0491L). PCR products were then purified using 0.7x to 1x AMPure XP Reagent (Beckman Coulter, A63881) depending on amplicon size and analyzed by automated electrophoresis using 4200 TapeStation System (Agilent, G2991BA). PCR Sequencing was performed by Plasmidsaurus Inc. using Oxford Nanopore Technologies followed by custom analysis and annotation.

Reads were mapped to the indicated reference genomes (Supplementary Table 2) using Minimap2 (V 2.22 --r1109dirty) ^2^, with specific alignment parameters optimized for SOS splicing events (minimap2 -ax splice -C 0) and hybrid spliceosome-SOS splicing events (minimap2 -ax splice). Isoform identification was performed using IsoQuant (V 3.5.0) ^3^ with parameters: isoquant.py --data_type assembly --check_canonical --keep_tmp --stranded none --report_canonical all --splice_correction_strategy none --model_construction_strategy sensitive_ont --report_novel_unspliced false. Nanopore sequencing coverage for specified genomic regions was normalized to the highest value, and splicing isoforms and splice sites were identified based on detected splicing events. Spliceosomal splice sites were identified from 10,000 randomly selected *C. elegans* introns using the annotation file WBcel235.gtf (Ensembl Release 104, WormBase WS276). Results were visualized using R (V 4.4.2).

### Automated electrophoresis using TapeStation System

Total RNAs were extracted from animals using the RNA Clean & Concentrator-5 Kit (Zymo Research, R1014) and reverse transcribed with Induro^®^ Reverse Transcriptase (NEB, M0681L) using Random Primer Mix, as described above. For PCR amplification, 50-200 ng of total DNA (to detect transposons in genomic DNA) or 25 ng of cDNA (to detect transposons in mRNA) was used as templates. Amplification was performed using Q5^®^ High-Fidelity DNA Polymerase (NEB, M0491L) for 30 cycles. PCR amplicons were analyzed using the 4200 TapeStation System (Agilent, G2991BA) with D1000 ScreenTape (Agilent, 5067-5582) or D5000 ScreenTape (Agilent, 5067-5588) following manufacturer’s instructions.

### SOS splicing reporter microinjection

The *Tc1::NmScarlet* SOS splicing reporter plasmid (Prpl-28::Tc1::NmScarlet::unc-54 3’ UTR) or variant constructs were microinjected at 50 ng/μL, along with 2 ng/μL of the co-injection marker pCFJ421 plasmid (Pmyo-2::gfp::h2b; Addgene, 34876). Pharyngeal GFP expression (co-injection marker) was used to detect successful microinjection. P0 animals were microinjected into the germline and allowed to lay a brood at 20 °C. F1 animals with pharyngeal GFP expression were scored for mScarlet expression.

### EMS screening and whole genome sequencing

*Tc1::rsd-3 (*i.e. *pk2013)*;*Tc1::Ngfp*-containing L4 stage animals were washed twice with M9 buffer and then mutagenized with 47 mM Ethyl methanesulfonate (EMS) by rotating at 20 °C for 4 hours. Mutagenized animals were grown on 100 mm NGM plates seeded with *E. coli* OP50 until most F1 animals had reached the young adult stage. F2 embryos were obtained via egg preparation from gravid adult hermaphrodites and placed on RNAi plates seeded with *E. coli* HT115 bacteria expressing *utp-20* dsRNA. Animals that failed to respond to *utp-20* RNAi (developed to young adults) were transferred to NGM plates seeded with *E. coli* OP50. Lineages established from these animals were tested for GFP expression. All potential mutants resistant to *utp-20* RNAi and failing to express GFP were further examined for reporter SOS splicing patterns using RT-PCR as described above.

A mapping strain was established from an animal mutagenized with EMS but that exhibited a wild-type response to *utp-20* RNAi (larval arrest) and normal GFP expression. To map mutants, males from the mapping strain were crossed with mutant hermaphrodites and eggs from F1 adult animals were collected and grown on *utp-20* RNAi plates. F2 animals that failed to respond to *utp-20* RNAi were scored for GFP expression. F2 animal lineages exhibited SOS splicing defects were pooled (>20 F2 animal lineages/mutant) in nearly equal proportions and subjected to genomic DNA extraction and whole genome sequencing at the Biopolymers Facility at Harvard Medical School or BGI Genomics. Reads were aligned to *C. elegans* genome (WBcel235/ce11) using BWA-MEM (V 0.7.17-r1188), and variants were identified using Samtools (V 1.3.1) and bcftools (V 1.13). Mapping SNPs were genotyped using GATK (V 4.1.9.0), and unique mutations absent in the mapping strains were plotted using R (V 4.4.2).

### Microscopic Imaging of *C. elegans*

For live worm imaging, worms were collected from NGM plate and washed once with M9 buffer. To restrain worm movement, 2% low melting agarose (Invitrogen, 16520050) was mixed with animals and then seeded onto glass slides. After agarose solidified, imaging was performed using a Nikon Ti2 W1 Yokogawa spinning disk confocal microscope equipped with Plan Apo λ 20x/0.8 DIC I or Plan Apo λD 60x/1.42 Oil DIC objective lens.

### RNA Immunoprecipitation and qRT-PCR

Approximately 5,000 young adult animals were washed three times with PBS/TX (PBS containing 0.01% (v/v) Triton X-100) and crosslinked with 1.8% formaldehyde by rotating at room temperature (RT) for 30 minutes. The reaction was neutralized by adding 125 mM glycine and rotated for 5 minutes at room temperature (RT). Crosslinked animals were washed with PBS/TX and resuspended in Pierce^TM^ IP Lysis/Wash Buffer (Thermo Scientific, 1861603) containing 80 U/mL RNaseOUT and protease inhibitor cocktail without EDTA (Roche, 11836170001) and sonicated (3 seconds on, 10 seconds off, 30% output for 2 minutes, 2 cycles. Lysates were rotated for an additional 15 minutes and clarified by centrifuging at 14,000 rpm for 15 minutes at 4 °C. The concentration of the supernatant was determined by Pierce™ BCA Protein Assay Kits (Thermo Scientific, 23225) and equal amounts of proteins from each sample were used for IP. Anti-FLAG^®^ M2 magnetic beads (used for 3xFLAG::AKAP-17; Millipore Sigma, M8823) were pre-blocked by 1% BSA with rotating 1 hour at RT and then incubated with supernatants at 4 °C overnight. Beads were washed with Pierce^TM^ IP Lysis Buffer for 6 times. Associated RNAs were reverse crosslinked by incubating at 70 °C for 1 hour and extracted with RNA Clean & Concentrator-5 (Zymo Research, R1014). cDNA was synthesized from isolated RNA using SuperScript™ IV Reverse Transcriptase (Invitrogen, 18090010). 5 ng of input cDNA per reaction was used. iTaq^TM^ Universal SYBR Green Supermix (Bio-Rad, 1725120) was employed for qRT-PCR analysis. Fold changes were normalized to inputs. All primers used in this study are listed in Supplementary Table 1.

### CRISPR

#### C. elegans

All CRISPR deletion or insertion experiments were done using CRISPR targeted genome editing techniques ^4^. crRNAs were designed using the IDT online guide RNA design tool (https://www.idtdna.com/site/order/designtool/index/CRISPR_SEQUENCE). For oligo-mediated Homology-Directed Repair (HDR), RNP complexes containing gene-specific crRNA (Supplementary Table 1), tracrRNA (IDT, 1072532), and Alt-R™ S.p.HiFi Cas9 Nuclease V3 (IDT, 1081060) were assembled at 37 °C, then mixed with ssODN (IDT, standard desalting; 4 nmol Ultramer) and a co-injection marker PRF4::rol-6(su1006) following manufacturer’s protocols. For dsDNA-mediated HDR (*akap-17::mScarlet*, *caap-1::mScarlet*), repair templates contained 35 bp homologous arms were amplified with PCR, gel-purified, cleaned with 1x to 1.5x AMPure XP Reagent (Beckman Coulter, A63881). Diluted repair template (100 ng/μL) was melted (95 °C 2 minutes, 85 °C 10 seconds, 75 °C 10 seconds, 65 °C 10 seconds, 55 °C 1 minute, 45 °C 30 seconds, 35 °C 10 seconds, 25 °C 10 seconds, 4 °C 10 seconds, with a 1 °C/s ramp down at each step) and mixed with RNPs before microinjection. Injection mix was microinjected into gonads of P0 animals and maintained at 20 °C. Rolling animals, which indicate successful injection, were isolated 4 days later and screened for successful genome deletion or insertion.

For integration of the *Tc1::Ngfp* SOS splicing reporter (Peft-3::Tc1::Ngfp::unc-54 3’ UTR), individual fragments were fused to generate the final genetic constructs using SapTrap cloning method as described ^5^. Engineered transgenes were subcloned into the pDD379 vector and integrated into the genome using CRISPR/Cas9.

#### Generation of HEK293T Knockout Cell Lines

To generate *AKAP17A* and *CAAP1* knockout (KO) cell lines in HEK293T cells, guide RNA (gRNA) sequences targeting *AKAP17A or CAAP1* were designed using CRISPick tool ^6^. Complementary DNA oligos encoding gRNA sequences (Supplementary Table 1) were synthesized from IDT, annealed and cloned into BsmBI-digested pLentiCRISPR-v2-BFP backbone ^7,8^ using Quick Ligation kit (NEB, M2200S). HEK293T cells were transfected with plasmids encoding gRNAs using Lipofectamine™ 3000 Transfection Reagent (Invitrogen, L3000008). Forty-eight hours post-transfection, single BFP-positive cells were isolated by Fluorescence-activated cell sorting (FACS) and cultured for 10-14 days for clonal expansion. Clonal populations were screened for knockout validation by immunoblot, confirming absence of protein expression from targeted genes.

#### Generation of SOS reporter Knock-in Cell Lines

To generate HEK293T cells expressing *Tc1-GFP* or *HSMAR2-GFP* reporters, SOS splicing reporters were cloned into donor plasmid AAVS1-tdTomato targeting vector (Addgene, 194728) using NEBuilder^®^ HiFi DNA assembly Master Mix (NEB, E2621S) with the following modifications; 1) tdTomato was replaced by Tc1-GFP or HSMAR2-GFP reporter cassettes; 2) PuroR selection marker was replaced with BlastR selection marker. Donor plasmids AAVS1-BlastR-Tc1-GFP or AAVS1-BlastR-HSMAR2-GFP, along with a Cas9 expressing plasmid targeting AAVS1 insertion site (pX458-AAVS1-sg; Addgene, 194721), were co-transfected into HEK293T cells using Lipofectamine™ 3000 Transfection Reagent (Invitrogen, L3000008). Three days post-transfection, 15 μg/mL Blasticidin HCl (InvivoGen, ant-bl-05) was added to select for edited cells. After seven days of selection, surviving cells were collected, and single GFP-positive cells were isolated in a 96-well plate by FACS. Resulting single-cell colonies were expanded, and knock-in cell lines were validated by Sanger sequencing and confirmed by observation of GFP expression.

### Lentivirus production and transduction

For lentiviral packaging, HEK293T cells cultured in plain DMEM were transfected with 2.5: 1: 1.5 ratio of the transfer plasmid pHAGE-2xFLAG-AKAK17A, VSV-G envelope-expressing plasmid pMD2.G (Addgene, 12259), and lentiviral packaging psPAX2 (Addgene, 12260) using CalPhos™ Mammalian Transfection Kit (Takara, 631312). After 12 hours, media was replaced with DMEM containing 10% FBS and the cells were incubated for 48 hours to produce lentiviral particles. Virus-containing supernatant was collected at 24-, 36-, and 48-hours post-transfection and filtered through a 0.45 μm syringe filter. Lentiviral particles were used to transduce he target cells with 10 μg/mL Polybrene Transfection Reagent (Sigma Aldrich, TR-1003-G). Following transduction, cells were selected with 2 μg/mL Puromycin Dihydrochloride (Gibco, A1113803) for 14 days. Puromycin-resistant cells were pooled for downstream analysis.

### Flow cytometry

HEK293T cells were rinsed once with PBS and detached from plates using TrypLE™ Express Enzyme (Gibco, 12605028), passed through a 35 μm nylon mesh strainer (Corning, 352235). Approximately 10,000 individual cells were analyzed for B525-FITC-A (green) using a CytoFLEX S flow cytometer. Flow cytometry data were collected using CytExpert software (Beckman Coulter), and figures were created using FlowJo (V 10.7.1) software. A Sony MA900 was used for fluorescence-activated cell sorting (FACS) to isolate isogenic single clones.

### SDS-PAGE and Immunoblotting

For cell lysate preparation, HEK293T cells were washed once with ice-cold PBS and lysed on ice for 5 minutes in RIPA Lysis and Extraction Buffer (Thermo Scientific, 89900) supplemented with PhosSTOP (Roche, 4906845001). Lysates were clarified by centrifugation at 20,000 g for 10 minutes at 4 °C. The supernatants were mixed with 4x SDS sample buffer (Millipore, 70607-3) supplemented with 4% β-mercaptoethanol (Sigma, M6250), denatured at 95 °C for 15 minutes and resolved using NuPAGE™ Bis-Tris Mini Protein Gels (Invitrogen, NP0322B0X) in 1x NuPAGE™ MOPS SDS Running Buffer (Invitrogen, NP0001).

After SDS-PAGE, proteins were transferred to nitrocellulose membranes (Bio-Rad, 1620112) using a semi-dry transfer method with a Power Blotter station (Invitrogen, PB0010). Membranes were blocked for 1 hour at room temperature with Intercept^®^ (TBS) Blocking Buffer (LI-COR Bio, 927-60001). Primary antibody (Supplementary Table 1) incubation was carried out at 4 °C overnight in TrueBlack^®^ WB Antibody Diluent (BIOTIUM, 23013B-1L). After washing with TBS-T (0.1% Tween 20), membranes were incubated with IRDye^®^ secondary antibodies (LI-COR Bio; Supplementary Table 1) for 1 hour. Membranes were washed with TBS-T and imaged using an Odyssey DLx Imaging System (LI-COR Bio).

### Immunoprecipitation and Mass Spectrometry

5 × 10^8^ HEK293T cells were seeded one day prior to transfection. HEK293T cells were transfected with plasmid expressing HA-CAAP1 (pGCS-3xHA-CAAP1) using CalPhos™ Mammalian Transfection Kit (Takara, 631312). After 24 hours, cells were lysed in 5 mL lysis buffer (40 mM HEPES pH 7.4, 100 mM NaCl, 0.05% CHAPS) with PhosSTOP (Roche, 4906845001) and sonicated as described above. Cell lysates were further rotated for 20 minutes at 4 °C and cleared by centrifugation at 21,000 g for 12 minutes. Supernatants were incubated with Pierce^TM^ Anti-HA Magnetic Beads (Thermo Scientific, 88836) at 4 °C for 2 hours on a rotator to immunoprecipitate HA-tagged CAAP1 protein complexes. Beads were washed four times with lysis buffer, and proteins were eluted using elution buffer (50 mM Tris-HCl pH 7.5, 10% SDS) by boiling at 95 °C for 4 minutes.

The eluted proteins were digested with Sequencing Grade Modified Trypsin (Promega, V5113) on S-Trap Micro columns (Protifi, C02-micro-10) according to manufacturer’s instructions. First, proteins were reduced using 5 mM Tris (2-carboxyethyl) phosphine hydrochloride (TCEP) (Sigma Aldrich, C4706-2G) at 55 °C for 15 minutes, followed by alkylation with 20 mM iodoacetamide (Sigma Aldrich, I6125) at room temperature in the dark for 30 minutes. After alkylation, samples were acidified with phosphoric acid (Sigma Aldrich, 345245-100 mL) to a final concentration of 2.5% (v/v). To assist in protein trapping, 10 volumes of 100 mM Tris (pH 7.55) in 90% methanol/10% water (v/v) were added, and the solution was passed through the S-Trap column by centrifugation at 4,000 g for 30 seconds. Multiple centrifugation rounds were performed to ensure columns were fully loaded. Once proteins were trapped, the column was washed three times with 100 mM Tris (pH 7.55) in 90% methanol/10% water (v/v) and spun dry. Proteins were digested by adding 2 µg of trypsin in 20 μL of 50 mM ammonium bicarbonate (pH 8) (Sigma Aldrich, A6141-25G). Digestion occurred overnight at 37 °C in a humidified environment. After digestion, peptides were eluted from the column in three steps, each by centrifugation at 4,000 g for 1 minute: 40 μL ammonium bicarbonate (pH 8), 40 μL 0.2% formic acid in water, and 40 μL 50% acetonitrile (Sigma Aldrich, 34851) in water. Eluted peptides were pooled, dried under reduced pressure using a SpeedVac (Eppendorf, 22820109), and re-suspended in 30 μL of 0.1% formic acid in water. LC-MS/MS data were acquired as previously described ^9^.

A protein database from the Human UniProt SwissProt proteome was used to identify proteins that co-immunoprecipitated with 3xHA-CAAP1. The FragPipe graphical user interface (V 18.0) was utilized to search data with MSFragger search engine and to post-process results. Tryptic peptides with up to two missed cleavages were included. Carbamidomethylation of cysteine was set as a fixed modification, and oxidation of methionine was allowed as a variable modification, with a maximum of four variable modifications per peptide. The allowed mass tolerances were 10 ppm for precursor ions and 0.04 Da for product ions. Peptide hits were filtered to a 1% false discovery rate using PeptideProphet in FragPipe.

### Co-immunoprecipitation

#### C. elegans Co-IP

Approximately 5,000 young adult animals were collected per sample and resuspended in Pierce^TM^ IP Lysis/Wash Buffer (Thermo Scientific, 1861603) containing protease inhibitor cocktail without EDTA (Roche, 11836170001) and sonicated using probe sonicator (10 seconds on, 10 seconds off, 50% output for 2 minutes with probe sonication). Lysates were rotated for 20 minutes at 4 °C and cleared by centrifugation at 21,000 g for 10 minutes. Protein concentration of supernatant was determined by the BCA method. For each Co-IP experiment, equal amounts of proteins from each sample were incubated with Myc-Trap Magnetic Agarose (ChromoTek, ytma-20) for 2 hours at 4 °C. Beads were washed four times with Pierce^TM^ IP Lysis/Wash Buffer (Thermo Scientific, 1861603). Proteins were eluted by adding 1x SDS sample buffer (Millipore, 70607-3) supplemented with 1% 2-mercaptoethanol (Sigma, M6250), followed by boiling at 95 °C for 15 minutes. Input and eluted proteins were separated with SDS-PAGE and detected with immunoblotting as described above.

#### Cell-based Co-IP

Co-IP experiments were performed using 2 × 10^6^ HEK293T cells, seeded in 60 mm dishes one day prior to transfection. Cells were transfected with the required plasmids using Lipofectamine™ 3000 Transfection Reagent (Invitrogen, L3000008). Forty-eight hours post-transfection, whole-cell lysates were prepared by adding 600 μL nuclear lysis buffer (1x PBS, 300 mM NaCl, 1% Triton X-100, 0.1% Tween 20) supplemented with PhosSTOP (Roche, 4906845001). Lysates were cleared by centrifugation at 20,000 g for 10 minutes at 4 °C and then incubated with Anti-FLAG^®^ M2 Magnetic Beads (for FLAG-AKAP17A; Millipore Sigma, M8823), Pierce^TM^ Anti-HA Magnetic Beads (for HA-RTCB; Thermo Scientific, 88836). Immunoprecipitation was performed by incubating lysates with beads on a tube rotator at 4 °C for 2 hours. Beads were washed three times with nuclear lysis buffer, and once with nuclear dialysis buffer (50 mM Tris-HCl pH 7.4, 100 mM NaCl, 0.1% Tween 20). Proteins were eluted by adding 1x SDS sample buffer, followed by boiling at 95 °C for 15 minutes. Both input and elution fractions were collected and analyzed by immunoblotting.

To capture RTCB-CAAP1 and RTCB-AKAP17A interactions, dithiobis (succinimidyl propionate) (DSP) crosslinking was performed as previously described ^10^. Before cell lysis, cells were washed twice with PBS, and protein complexes were crosslinked using 0.1 mM DSP (Thermo Scientific, 22586) in PBS at 37 °C for 30 minutes. Crosslinking reactions were quenched by adding 20 mM Tris-HCl in PBS (pH 7.4) for 15 minutes at room temperature. Cells were lysed and proceed with Co-IP protocol described above.

### Tripartite splitGFP reporter

To monitor protein-protein interactions for AKAP17A, CAAP1, and RTCB, 5 × 10^4^ HEK293T WT or *CAAP1* knockout cells were seeded into 24-well glass-bottom plate (Cellvis, P24-1.5H-N) one day prior to transfection. The next day, cells were co-transfected with plasmids encoding for: GFP1-9 (Addgene, 182244), GFP10 (pGCS-HA-GFP10), GFP11 (pGCS-FLAG-GFP11) (negative control); AKAP17A-GFP10, GFP10, and GFP1-9 (negative control for AKAP17A); CAAP1-GFP10, GFP11, and GFP1-9 (negative control for CAAP1); RTCB-GFP11, GFP10, GFP1-9 (negative control for RTCB); AKAP17A-GFP10, RTCB-GFP11, and GFP1-9 (AKAP17A and RTCB interaction); CAAP1-GFP10, AKAP17A-GFP11, and GFP1-9 (CAAP1 and AKAP17A interaction); CAAP1-GFP10, RTCB-GFP11, and GFP1-9 (CAAP1 and RTCB interaction) using Lipofectamine™ 3000 Transfection Reagent. Twenty-four hours after transfection, cells were washed with PBS and the culture media was replaced with live-cell imaging media. Imaging was performed 30 minutes after media exchange using a Nikon Ti2 W1 Yokogawa spinning disk confocal microscope equipped with Plan Apo λD 60x/1.42 Oil DIC objective lens.

### Illumina sequencing and alternative splicing analysis

Total RNA was extracted using TRIzol^TM^ Reagent (Invitrogen, 15596026), followed by DNase I (Thermo Scientific, EN0521) treatment to remove contaminating DNA. RNA concentrations were determined with Nanodrop^TM^ 2000, and RNA quality was assessed via gel electrophoresis. Ribosomal RNA and mitochondrial RNA were depleted using fifty nucleotide DNA oligos complementary to *C. elegans* rRNA and mtRNA sequences, followed by Thermostable RNase H (NEB, M0523S) treatment, as described in ^11^. Ribosomal-depleted RNA was then analyzed using TapeStation and qPCR, following quality control methods outlined in manufacturer instructions. Libraries were prepared using the KAPA mRNA HyperPrep Kit (KR1352-v4.17) and sequenced with 150 nt paired-end reads, generating 30 million read pairs on the Illumina NovaSeq 6000 (Biopolymers Facility, Harvard Medical School).

Reads were processed using the Nextflow (V 24.04.4)-based nf-core/rnasplice pipeline ^12^ (V 1.0.4), primarily for downstream rMATS analysis ^13^ (V 4.1.2). Read quality was assessed using FastQC (https://www.bioinformatics.babraham.ac.uk/projects/fastqc/) (V 0.12.1), and low-quality reads were trimmed or removed using Trim Galore (https://www.bioinformatics.babraham.ac.uk/projects/trim_galore/) (V 0.6.7). Adapter sequences and low-quality bases were clipped. Remaining reads were aligned to the C. elegans genome (WBcel235/ce11) using STAR 14 (V 2.7.9a). The resulting alignment was subjected to rMATS for annotation of alternative splicing events. Differential alternative splicing events were identified with cutoff criteria: ΔPSI ≥ 0.05 or ≤ −0.05, *P*-value < 0.05. Results were visualized using boxplots.

### *In vitro* protein synthesis and pull-down

HA::RTCB-1 protein was synthesized using the PURExpress ^®^ *In Vitro* Protein Synthesis Kit (NEB, E6800S) for *C. elegans* HA::RTCB-1 pull-down experiments. *C. elegans rtcb-1* was codon-optimized for *E. coli* expression and synthesized by Twist Bioscience. The codon-optimized *rtcb-1* sequence was cloned into the pDFCI vector with an N-terminal 3×HA tag. Protein synthesis reactions were performed according to manufacturer’s instructions. Following protein synthesis, the reaction was stopped by dilution in 500 μL of Pierce^TM^ IP Lysis/Wash Buffer (Thermo Scientific, 1861603) and incubated overnight at 4 °C with Pierce^TM^ Anti-HA Magnetic Beads (Thermo Scientific, 88836) to capture HA::RTCB-1.

For pull-down, worm lysates were prepared as described above and incubated with HA::RTCB-1-bound beads at 4 °C for 2 hours. Beads were washed four times with Pierce^TM^ IP Lysis/Wash Buffer, and bound proteins were eluted using 1x SDS sample buffer. Both input and elution fractions were collected and analyzed by immunoblotting.

### AlphaFold3 Multimer prediction

For each AlphaFold3 protein-protein interaction and docking prediction, the full sequence of proteins was used as input ^15^. AlphaFold3-multimer prediction was performed on AlphaFold Server (https://alphafoldserver.com) to generate 4 predicted unrelaxed docking structures with default parameters. For each docking prediction, the highest scoring predicted structure is shown and illustrated using Pymol (V 2.5.8).

## Main text statement

## Code availability

Descriptions of custom scripts used to analyze Nanopore long-read sequencing and Illumina RNA sequencing data are provided in the Methods section. The scripts are available upon request from the corresponding author.

## Acknowledgements

We thank members of the Kennedy laboratory, particularly David D. Lowe for helpful discussions and Dr. Cindy Chang for sharing the *Tc1::Ngfp* strain, Dr. David Wassarman for helpful conservations, Dr. Steven Gygi and the Taplin Mass Spectrometry Facility at HMS for the use of their mass spectrometers and the Biopolymers Facility at Harvard Medical School (HMS) for Illumina sequencing. We are grateful to Dr. Wells Adrienne and Dr. Praju Vikas Anekal from the Microscopy Resources On The North Quad (MicRoN) Facility for microscopy assistance, as well as Alex Bartlett, Dr. Lorena Pantano, and Dr. Shannan Ho Sui from the Harvard Chan Bioinformatics Core for discussions and for contributing scripts to RNA splicing analysis optimization. Some strains were provided by the Caenorhabditis Genetics Center (CGC) (P40 OD010440). This work was supported by NIH grants R35GM148206 (S.K.) and AG11085 (S.J.E.), and the Howard Hughes Medical Institute (HHMI).

## Author contributions

L.Z. designed and performed all experiments and generated all figures. C.N. and S.J.E. assisted with cell line work and contributed to Figs. 4d-e, 5a, g, and Extended Data Figs. 9d-e, 10g. J.A.P. assisted with Mass Spectrometry, contributed to Fig. 5a. S.K. assisted with the genetic screen and contributed to Figs. 3a-b. S.K. and L.Z. conceived the project. S.K. supervised the study and wrote the manuscript with edits from all authors.

## Ethics declarations

## Competing interests

S.J.E. is a founder of TSCAN Therapeutics, MAZE Therapeutics, ImmuneID, and Mirimus, serves on the scientific advisory boards of Homology Medicines, ImmuneID, MAZE Therapeutics, X-Chem, and TSCAN Therapeutics, and is an advisor for MPM Capital. Other authors declare no competing interests.

## Supplementary information

Supplementary Figure 1

This figure includes the uncropped gel images and Tapestation results.

Supplementary Table 1

Oligos, *C. elegans* strains, cell lines and antibodies used in this study.

Supplementary Table 2

Reference sequences used for SOS splicing analysis.

## Extended Data Figure legends

**Extended Data Figure 1.**
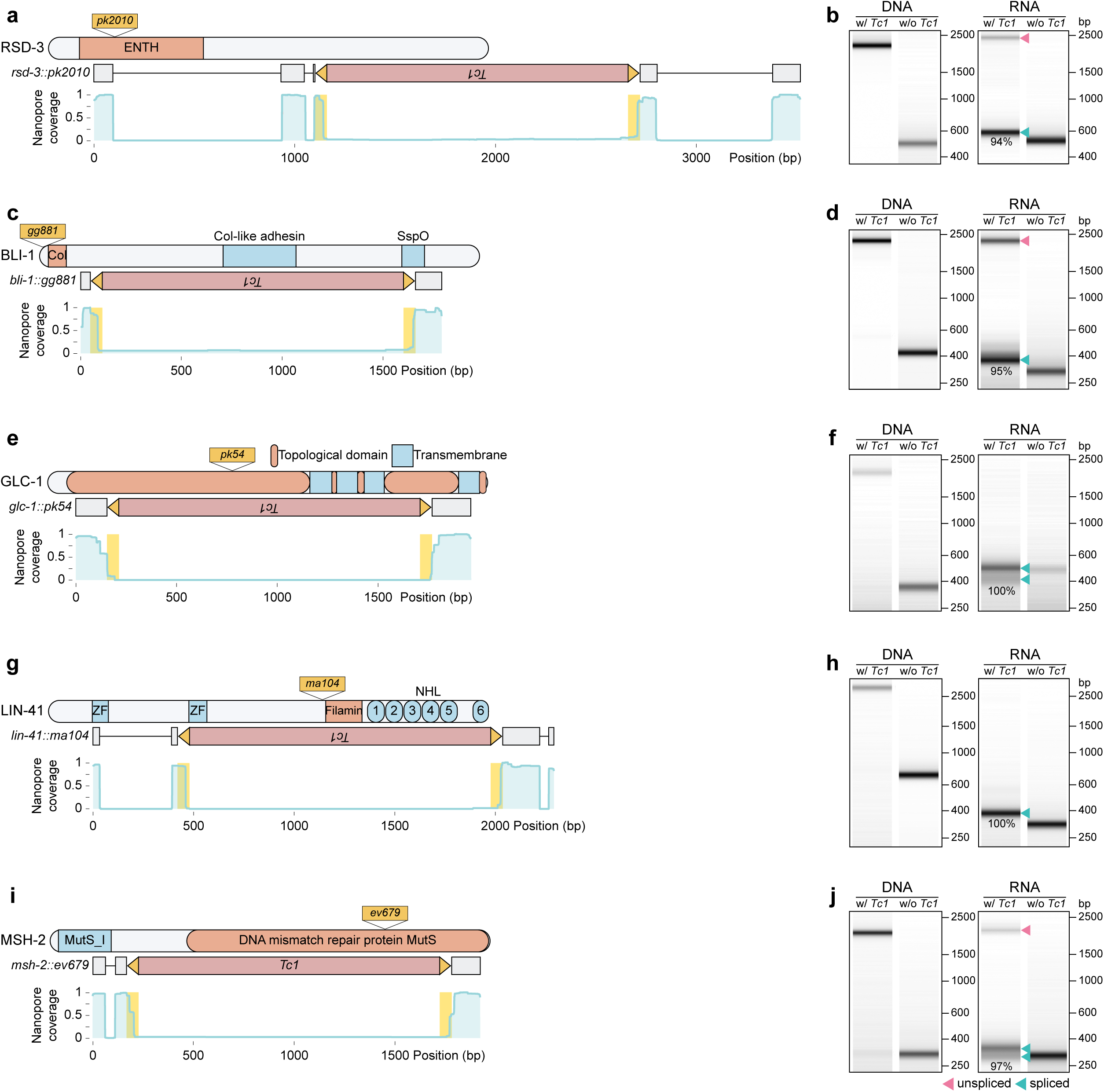
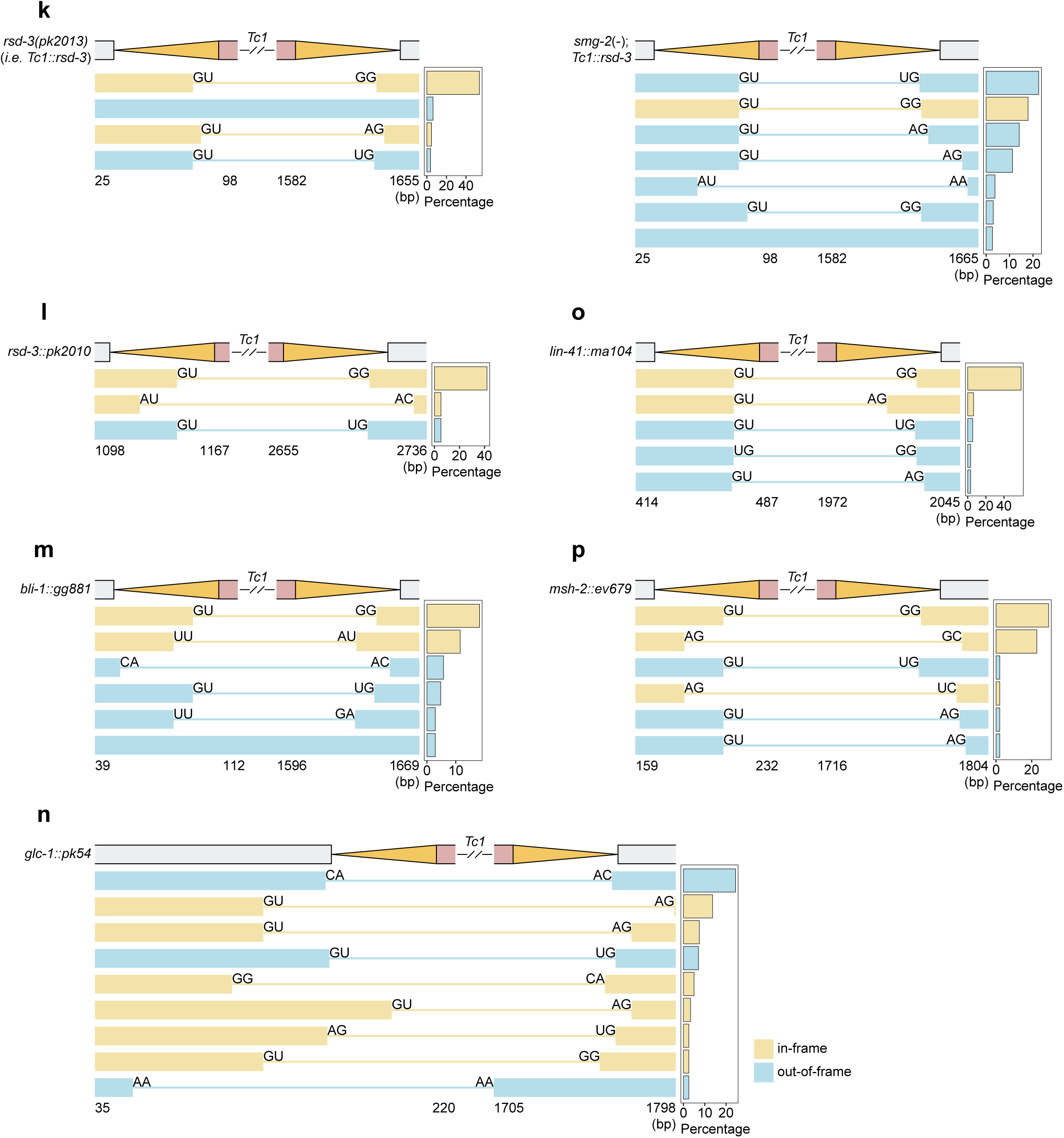
SOS splicing will remove any *Tc1* element from its host mRNA. (Page 1, Left) Nanopore sequencing of RNA isolated from *rsd-3::pk2010* (**a**), *bli-1::gg881* (**c**), *glc-1::pk54* (**e**), *lin-41::ma104* (**g**), *msh-2::*ev679 (**i**) animals. PCR amplicons were generated using primers flanking the *Tc1* transposon insertion site. Primer sites are indicated. *Tc1 ITR* regions are highlighted. Protein domain for each gene, and *Tc1* insertion sites, are also indicated. (Page 1, Right) SOS splicing visualized by Tapestation. PCR amplicons were generated from genomic DNA (left panel) or cDNA (RT-PCR) (right panel) isolated from *rsd-3::pk2010* (**b**), *bli-1::gg881* (**d**), *glc-1::pk54* (**f**), *lin-41::ma104* (**h**), *msh-2::*ev679 (**j**) animals (w/ *Tc1*). Wild-type (N2) animals (w/o *Tc1*) serve as negative controls. Unspliced and spliced amplicons are indicated. SOS splicing efficiency is indicated. (Page 2) SOS splicing isoforms detected in *rsd-3(pk2013)* (referred to as *Tc1::rsd-3* in main text) (**k**, left panel), *smg-2*(−);*rsd-3(pk2013)* (**k**, right panel), *rsd-3::pk2010* (**l**), *bli-1::gg881* (**m**), *glc-1::pk54* (**n**), *lin-41::ma104* (**o**), *msh-2::*ev679 (**p**), are shown and ranked by abundance. Isoforms representing > 2% of total reads are shown.

**Extended Data Figure 2.**
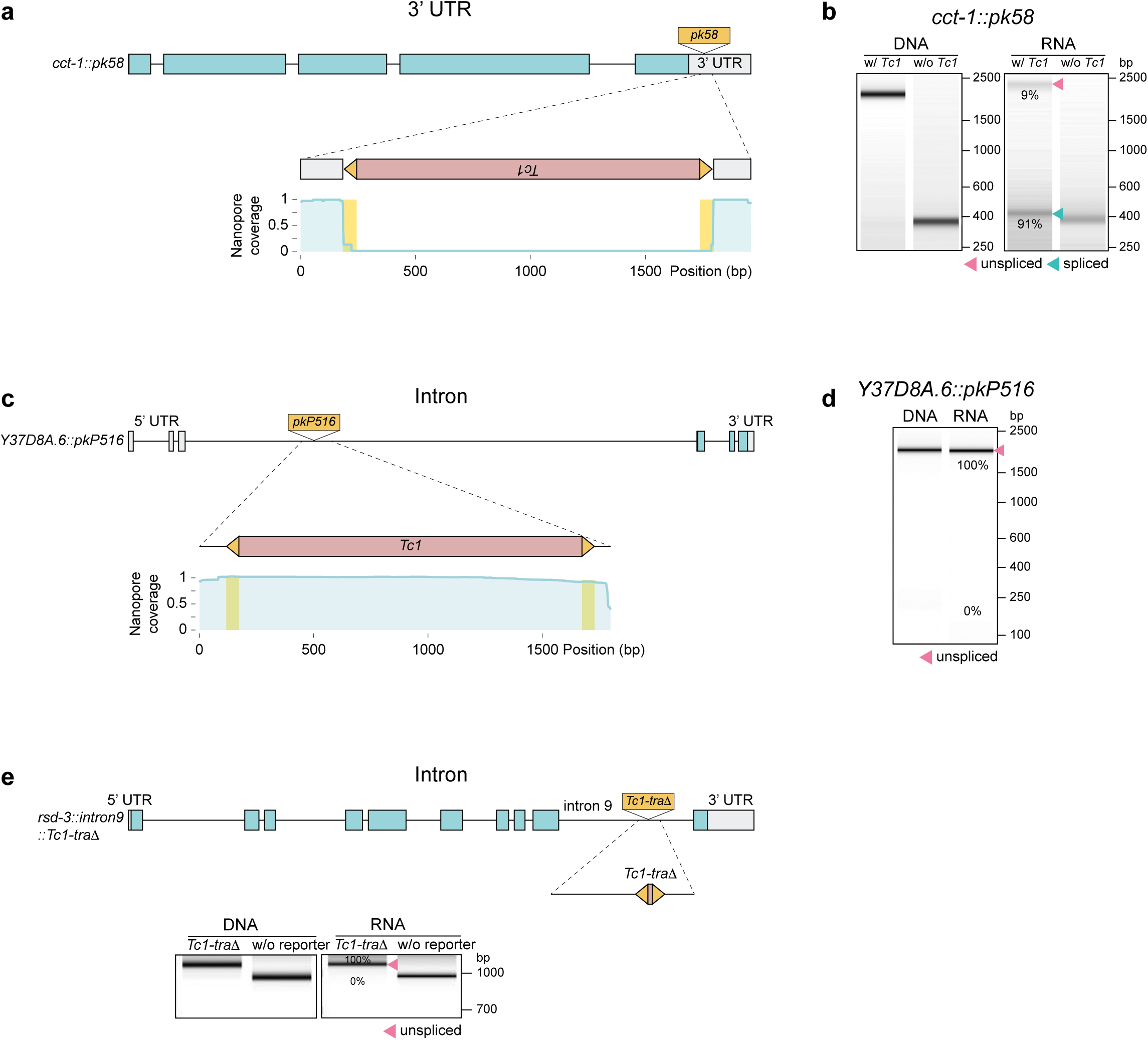
SOS splicing of *Tc1* from a 3’ UTR, but not from introns. **a, c.** Nanopore sequencing of RNA isolated from (**a**) *cct-1::pk58* (present in 3’ UTR) and (**c**) *Y37D8A.6::pkP516* animals (present in intron). (**b, d, e**). SOS splicing of (**b**) *cct-1::pk58*, (**d**) *Y37D8A.6::pkP516*, and (**e**) A *Tc1* derivative (*Tc1-traΔ),* which lacks 98% of the *Tc1* transposase gene but is, nonetheless, still subjected to SOS splicing when placed in exons (see Fig. 2) was inserted into the ninth intron of *rsd-3* and visualized by RT-PCR and Tapestation. PCR amplicons were generated from genomic DNA (left panel) or cDNA (RT-PCR) (right panel). Unspliced and spliced amplicons with percentage are indicated.

**Extended Data Figure 3.**
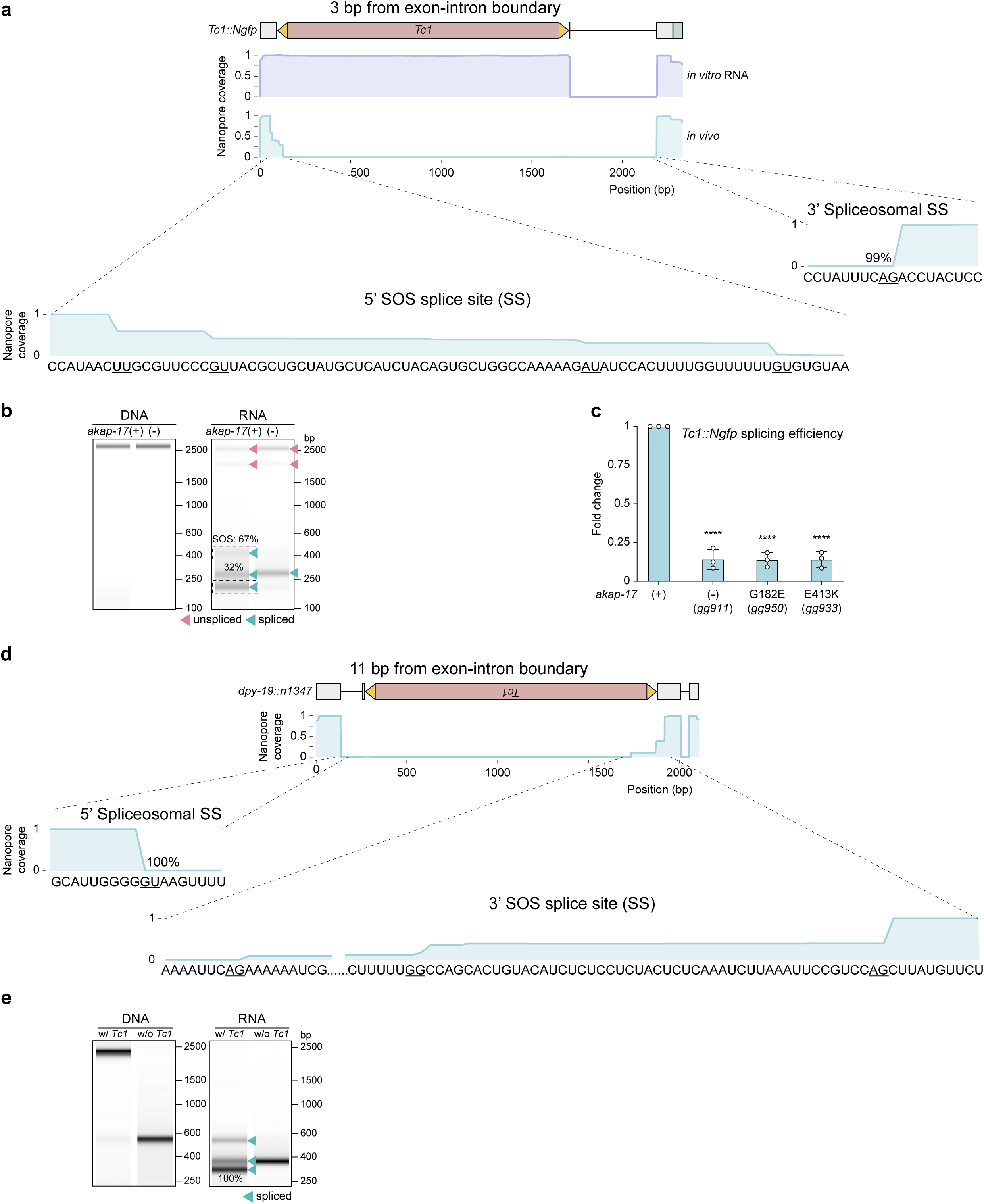
Hybrid spliceosome-SOS splicing. **a.** Nanopore sequencing of RNA isolated from *Tc1::Ngfp* animals. *In vitro* synthesized RNAs were included as controls. In *Tc1::Ngfp* animals, *Tc1* is located 3 nucleotides from a canonical 5’ host-gene spliceosome splice site. Sequence surrounding observed splice sites are shown, with common splice sites underlined. The data show that *Tc1* is excised from *Tc1::Ngfp* via 5’ splice sites in or near the 5’ *ITR* of *Tc1* and a canonical spliceosome 3’ splice site in the downstream host-gene exon. These splice isoforms, which connect apparent SOS splice sites (see Figure 1 of the main text) and canonical spliceosome splice sites, are referred to as hybrid SOS-spliceosome splice isoforms. The existence of these RNAs suggests that, remarkably, the two splicing systems (SOS and spliceosome) can, in some contexts, coordinate their splicing efforts. Assessing how and when the SOS and spliceosomal splicing systems interact could reveal additional biological function(s) of SOS splicing. Note that some of the *Tc1* excision events from *Tc1::Ngfp* are GU-AG splices (see below). **b.** Because *Tc1::Ngfp* is one of the two reporter genes used for the genetic screen presented in Figure 3 of the main text, a more detailed description of *Tc1* excision patterns from *Tc1::Ngfp* is presented in panels (**b-c**). *Tc1::Ngfp* splice isoforms visualized by Tapestation analysis are shown. PCR amplicons were generated from genomic DNA (left panel) or cDNA (RT-PCR) (right panel) isolated from animals of indicated genotypes. Data to be presented in Figure 3 of the main text will show that the *akap-17* gene product is required for 90-99% of SOS splicing events that occur at loci in which *Tc1* is integrated distal to host-gene splice sites, such as all the *Tc1* insertions analyzed in Figure 1 of the main text and ED Figure 1. The data in this panel (**b**) show that production of some (but not all, see below) hybrid SOS-spliceosome *Tc1::Ngfp* splice isoforms depend upon AKAP-17 for their production. These data support the idea that the splice isoforms we refer to in panel (**a**) as hybrid SOS-spliceosome splice isoforms above depend upon SOS splicing for their biogenesis, as expected. The AKAP-17 independent *Tc1::Ngfp* isoforms shown in panel (**b**) are surprising because they are not observed in sequencing of other *Tc1* insertion alleles and the provenance of these isoforms is not yet known. We note, however, that a subset of the *Tc1* excision isoforms shown in panel (**a**) are GU-AG splices, which could, due to the fact that they are GU-AG, be spliceosome derived. We speculate that when *Tc1* inserts very near (*e.g.* 3 nt) a 5’ splice site (or 3’ splice site, see panel (**d-e**)), insertion prevents the spliceosome from recognizing nearby spliceosome splice sites, causing the spliceosome to search for and identify distinct donor or acceptor splice sites. Why might *Tc1* happen to possess a canonical splice site-whose use enables *Tc1* excision via the spliceosome (AKAP-17-independent)-when spliceosome sites are disrupted by *Tc1* insertion? Transposons do not benefit when they harm their hosts. For this reason, transposons such as *Tc1* could be expected to evolve in ways that limit their interference with host gene expression. Given these considerations, we speculate that *Tc1*, and perhaps other transposons, have evolved consensus spliceosome 5’ splice sites (or 3’, see panel (**c**)) in their *ITR* elements, which can be used by host cells to excise TEs, via the spliceosome, when *Tc1* lands near and disrupts a host donor (or acceptor, see panel (**d-e**)) splice site. When taken together with all the other data presented in this work, the result in this panel lead us to the following model: *C. elegans* has both a primary and a backup system to remove TEs from host mRNAs; 1) SOS splicing, which is the primary pathway and the subject of this work, excises most TEs from mRNAs independently of the spliceosome; and 2) a back-up pathway that uses the spliceosome to remove TEs when they land near and disrupt host donor or acceptor splice site. *Tc1::Ngfp* was chosen to be one of the two reporter genes used in our screen before we understood the complexity of splicing at this locus. Future screens that use reporter genes that report solely on the production of hybrid SOS-spliceosome splicing could identify host factors that drive hybrid spliceosome-SOS splicing. **c.** qRT-PCR analysis quantifying efficiency of *Tc1* excision from *Tc1::Ngfp* in animals of the indicated genotypes. Splicing efficiency was determined by calculating the percentage of spliced mRNA relative to total *Tc1::Ngfp* reporter mRNA, with splicing efficiency of wild-type animals serving as a normalization standard (set to 1). *Tc1* excision from *Tc1::Ngfp* in three independent *akap-17* mutant strains is decreased 2-3x. The data support the idea that most, but not all, *Tc1* excision events from *Tc1::Ngfp* depend on the SOS splicing machinery, which is consistent with data in panel (**b**). We suspect that residual *Tc1* excision observed from *Tc1::Ngfp* in *akap-17(−)* animals represents GU-AG spliceosomal splicing, as outlined in panel (**b**). Data are presented as mean ± SD with all data points shown. *N* = 3 biologically independent experiments. One-way ANOVA analysis followed by Tukey’s post-hoc test was performed to assess statistical significance. \*\*\*\**P* < 0.0001. **d.** Nanopore sequencing of RNA isolated from *dpy-19::Tc1(n1347)* animals. In *dpy-19::Tc1* animals, *Tc1* is located 11 nucleotides from a canonical 3’ host-gene spliceosome splice acceptor site. Sequences bordering splice sites (SS) are shown, with common splice sites underlined. Hybrid SOS-spliceosome isoforms are detected in *dpy-19::Tc1* animals, which connect apparent SOS splice sites in or near the 3’ ITR of *Tc1* to a neighboring spliceosome donor site. The data also show that in *dpy-19::Tc1*(*n1347*) animals, similar to what was observed for the *Tc1::Ngfp* gene (**b**), about half of the splice isoforms generated by *dpy-19::Tc1* are GU-AG splices and about half are not. We have not yet tested the AKAP-17 (SOS splicing) dependence of these isoforms. We predict that in *akap-17*(−) animals, the non-GU-AG *dpy-19::Tc1* splice isoforms will be missing because these RNAs are hybrid SOS-spliceosome splices, while the GU-AG *dpy-19::Tc1* splice isoforms will persist, because they derive exclusively from the spliceosome. **e.** Splicing isoforms of *dpy-19::Tc1* visualized by RT-PCR and Tapestation analysis. Spliced amplicons with percentage are indicated.

**Extended Data Figure 4.**
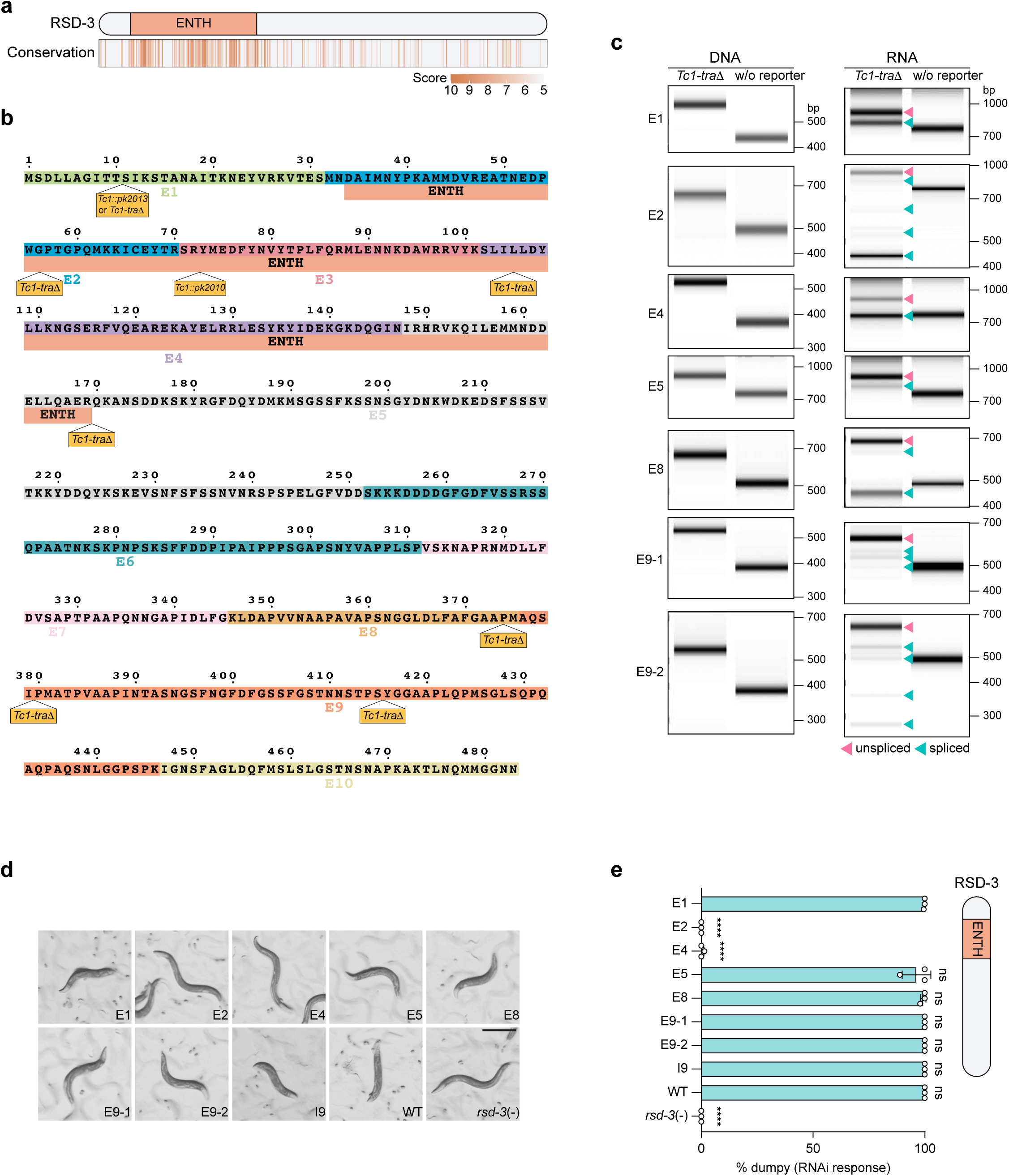
Position dependent rescue of gene function by SOS splicing. **a.** *C. elegans* RSD-3 protein and its orthologs sequences from *S. cerevisiae*, *A. thaliana*, *D. melanogaster*, *D. rerio*, *X. laevis*, *M. musculus*, and *H. sapiens* were aligned using Clustal Omega multiple sequence alignment program. Conservation scores generated from ConSurf analysis are represented by a color gradient. The ENTH domain is well-conserved across species. **b.** RSD-3 protein sequence showing locations of transposon insertions within the *rsd-3* gene. Full-length *Tc1* elements (*Tc1*), or a *Tc1* derivative (*Tc1-traΔ),* which lacks 98% of the *Tc1* transposase gene but is, nonetheless, subjected to SOS splicing when inserted into exons (see Fig. 2), were inserted into the indicated eight locations in *rsd-3*. Amino acids encoded by different exons are demarcated with different colors. The ENTH (epsin N-terminal homology) domain, which is the only portion of RSD-3 that is conserved in other animals, is highlighted. **c.** SOS splicing visualized by RT-PCR and Tapestation analysis. PCR amplicons were generated from genomic DNA (left panel) or cDNA (RT-PCR) (right panel) isolated from the specified *Tc1-traΔ* insertions. Wild-type (N2) animals (w/o reporter) serve as negative controls. Unspliced and spliced amplicons are indicated. **d.** DIC images of animals harboring the indicated *Tc1-traΔ* knock-in alleles from (**a**) and ED Fig. 2e, treated with *dpy-6* dsRNA. *rsd-3*(−) animals, which harbor a *rsd-3* deletion, serve as positive controls for loss of RSD-3 function. Scale bar = 200 μm. **e.** (Left) Quantification of RNAi responsiveness to *dpy-6* dsRNA. % of animals exhibiting dumpy (Dpy) phenotypes is indicated. (Right) Sites of *Tc1-traΔ* insertion relative to RSD-3, and the RSD-3 ENTH domain, are shown. Data are presented as mean ± SD with all data points shown. *N* = 3 biologically independent experiments. One-way ANOVA analysis followed by Tukey’s post-hoc test was performed to assess statistical significance. \*\*\*\**P* < 0.0001, ns not significant, *P* > 0.05.

**Extended Data Figure 5.**
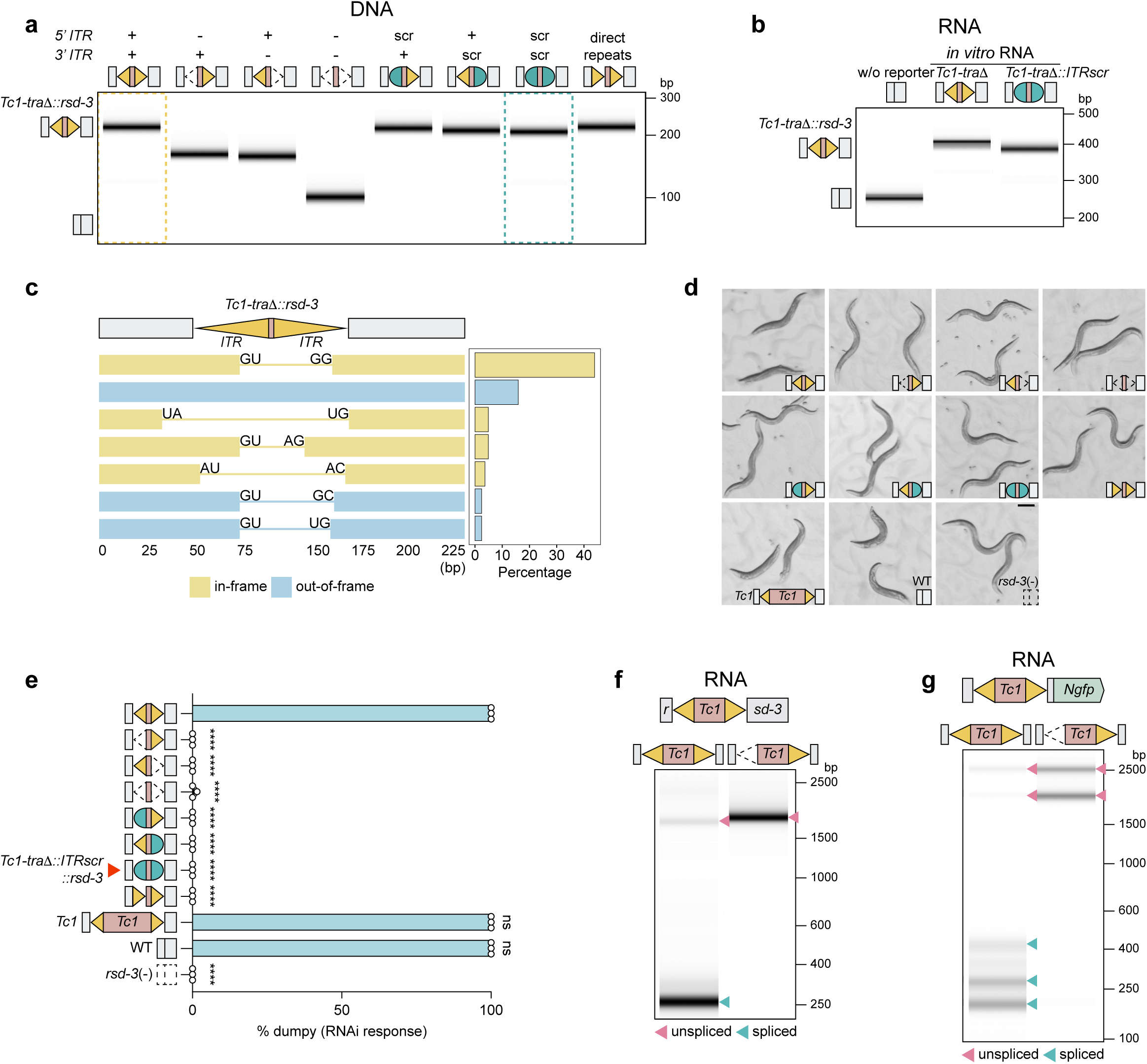
*ITR* base-pairing triggers SOS splicing. **a.** Tapestation analysis showing a lack of *Tc1-traΔ* excision from genomic DNA from animals harboring the indicated *Tc1-traΔ* variants. This data is a control for results shown in Fig. 2d in the main text. (Top) Schematic of variants tested. Wild-type (N2) animals (w/o *Tc1-traΔ*) serve as negative controls. Scr, scrambled *ITR*. **b.** *In vitro* transcribed *Tc1-traΔ* or *Tc1-traΔ::ITRscr* RNAs were analyzed by RT-PCR and Tapestation. The results show that SOS splicing does not occur during library preparation or sequencing for *Tc1-traΔ* or *Tc1-traΔ::ITRscr* RNAs, establishing that the SOS splicing events shown in Fig. 2d of the main text occur *in vivo*. Wild-type (N2) animals (w/o *Tc1-traΔ*) serve as negative controls. **c.** SOS splicing isoforms detected by nanopore sequencing in *Tc1-traΔ::rsd-3* mRNA, ranked by abundance. Isoforms representing > 2% of total reads are shown. **d.** DIC images of animals harboring the indicated *Tc1-traΔ* variant alleles treated with *dpy-6* dsRNA. (bottom right corner) Schematic of variants. Scale bar = 200 μm. **e.** Quantitation of RNAi responses to *dpy-6* dsRNA. % of animals exhibiting dumpy (Dpy) phenotypes is shown. Note that *Tc1-traΔ::ITRscr::rsd-3* animals were defective for RNAi (indicated by red arrow), which is not the expected result based upon our *ITR* base-pairing triggers SOS splicing model. Lack of RSD-3 rescue in *Tc1-traΔ::ITRscr* animals is not due to a failure of base-paired *ITRscr* to trigger SOS splicing in this RNA (see main text, Fig. 2e, f, which shows that SOS splicing does occur) but, rather, we suspect because of the relatively poor efficiency of SOS splicing for this mRNA (see main text, Fig. 2e) and because relatively few SOS splice events, which do occur in this RNA, happen to be in-frame (see main text, Fig. 2f). Data are mean ± SD with all data points shown. *N* = 3 biologically independent experiments. One-way ANOVA analysis followed by Tukey’s post-hoc test was performed to assess statistical significance comparing to animals with *Tc1-traΔ* allele. \*\*\*\**P* < 0.0001, ns not significant, *P* > 0.05. **f.** Tapestation-based detection of SOS splicing in RNA isolated from *Tc1::rsd-3* and *Tc1-5’ ITRΔ::rsd-3* animals. Unspliced and spliced amplicons are indicated. **g.** Tapestation-based detection of SOS splicing in RNA isolated from *Tc1::unc-54* and *Tc1-5’ ITRΔ::unc-54* animals. Unspliced and spliced amplicons are indicated.

**Extended Data Figure 6.**
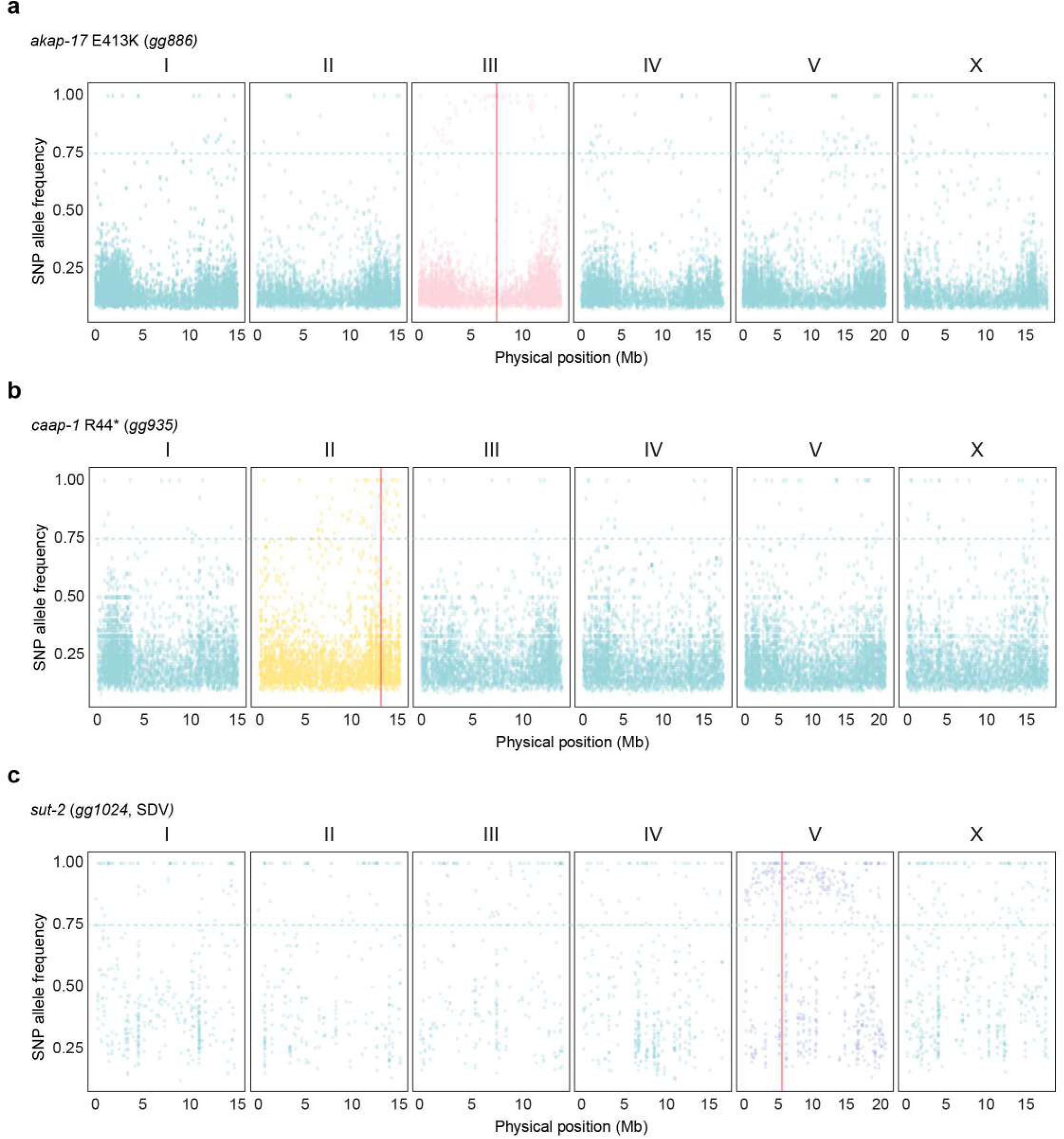
Positional mapping of SOS splicing mutations. Mutants were outcrossed and >20 F2 progeny exhibiting SOS splicing defects were isolated. Genomic DNA from pooled F2s was sequenced. Chromosomal regions linked to the SOS splicing genes should be enriched for EMS-based mutations in these F2 animals. Allele frequencies for EMS-induced mutations in F2 progeny of *gg886* (**a**), *gg935* (**b**), and *gg1024* (**c**) crosses are shown for the six *C. elegans* chromosomes. Red line denotes the location of the indicated candidate SOS splicing factor. Chromosome positions are shown on the x-axis (in Mb), and SNP allele frequencies are shown on the y-axis. The dotted line indicates EMS SNP allele frequency greater than 75%. SNP allele frequencies were calculated using the bcftools algorithm.

**Extended Data Figure 7.**
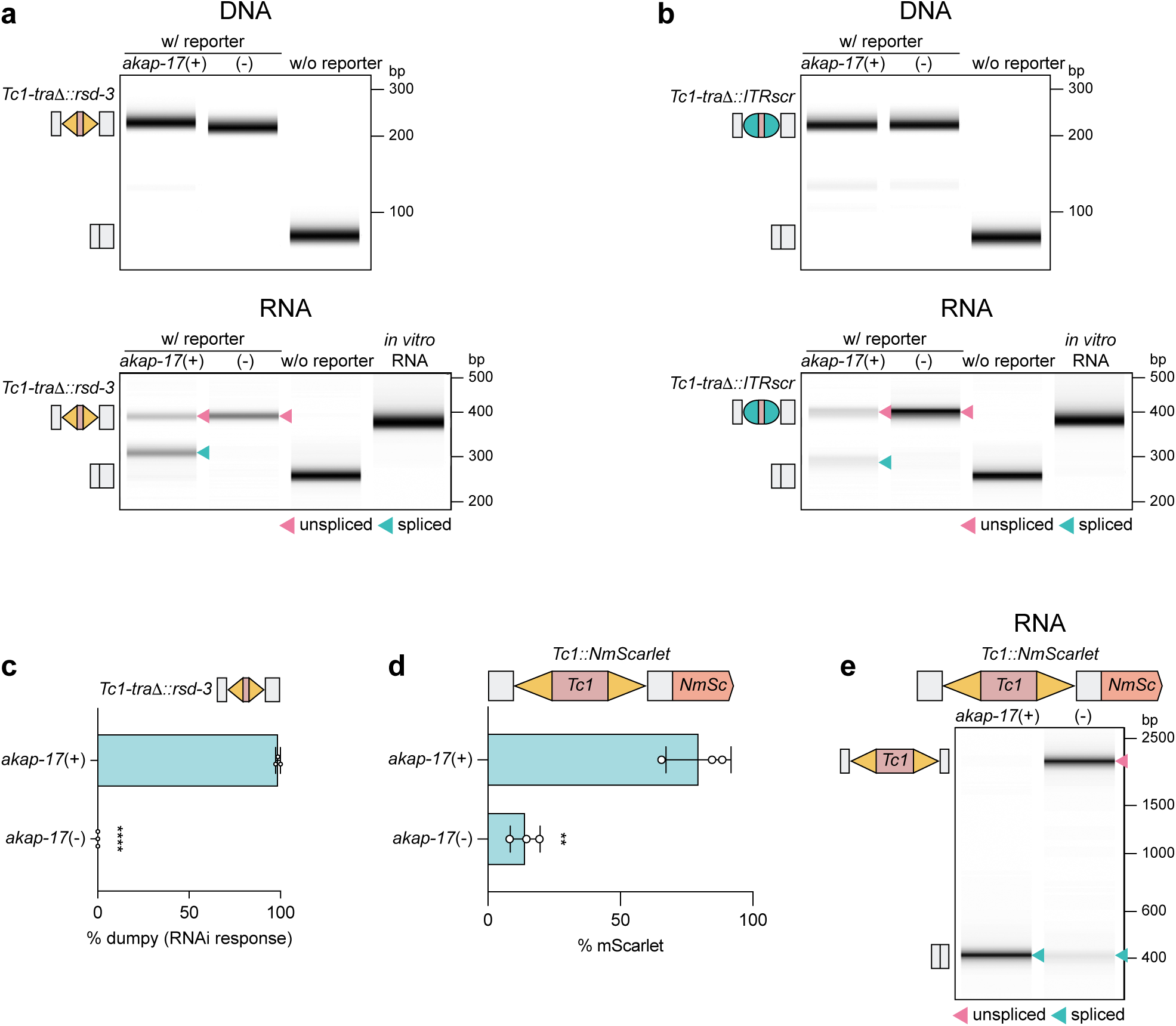
*akap-17*, *caap-1* and *sut-2* are required for SOS splicing. **a-b**. Tapestation analysis showing that SOS splicing of *Tc1-traΔ::rsd-3* and *Tc1-traΔ::ITRscr::rsd-3* RNAs requires AKAP-17. (Top) As expected, DNA from animals harboring (**a**) *Tc1-traΔ::rsd-3* or (**b**) *Tc1-traΔ::ITRscr::rsd-3* does not show *Tc1* excision. (Bottom) (**a**) RT-PCR followed by Tapestation analysis shows that the *Tc1-traΔ::rsd-3* and (**b**) *Tc1-traΔ::ITRscr::rsd-3* RNA are SOS spliced and that this splicing depends upon a wild-type copy of *akap-17*. For all panels in this figure, *akap-17*(−) is *akap-17(gg911)*. Wild-type (N2) animals (w/o *Tc1-traΔ*) serve as negative controls. Scr, scrambled *ITR*. Unspliced and spliced amplicons are indicated. **c.** *akap-17*(+) or (−) animals that express *Tc1-traΔ::rsd-3* were treated with *dpy-6* dsRNA. % of animals exhibiting dumpy (Dpy) phenotypes is indicated. The data show that AKAP-17 is required for rescuing RSD-3 functionality in *Tc1-traΔ::rsd-3* animals. Data are presented as mean ± SD with all data points shown. *N* = 3 biologically independent experiments. The two-tailed unpaired student’s *t*-test. \*\*\*\**P* < 0.0001. Scale bar = 200 μm. **d.** *Tc1::NmScarlet* SOS splicing reporter was injected into P0 adult animals of the indicated genotypes, and the percentage of F1 progeny expressing mScarlet signal was quantified. The data show that AKAP-17 is required for rescuing mScarlet expression from *Tc1::NmScarlet* SOS. Data are presented as mean ± SD with all data points shown. *N* = 3 biologically independent experiments. The two-tailed unpaired student’s *t*-test. \*\**P* < 0.01. **e.** Tapestation-based detection of *Tc1::NmScarlet* SOS splicing in F1 progeny animals of the indicated genotypes. The data show that AKAP-17 is required for SOS splicing of *Tc1::NmScarlet* SOS. Unspliced and spliced amplicons are indicated.

**Extended Data Figure 8.**
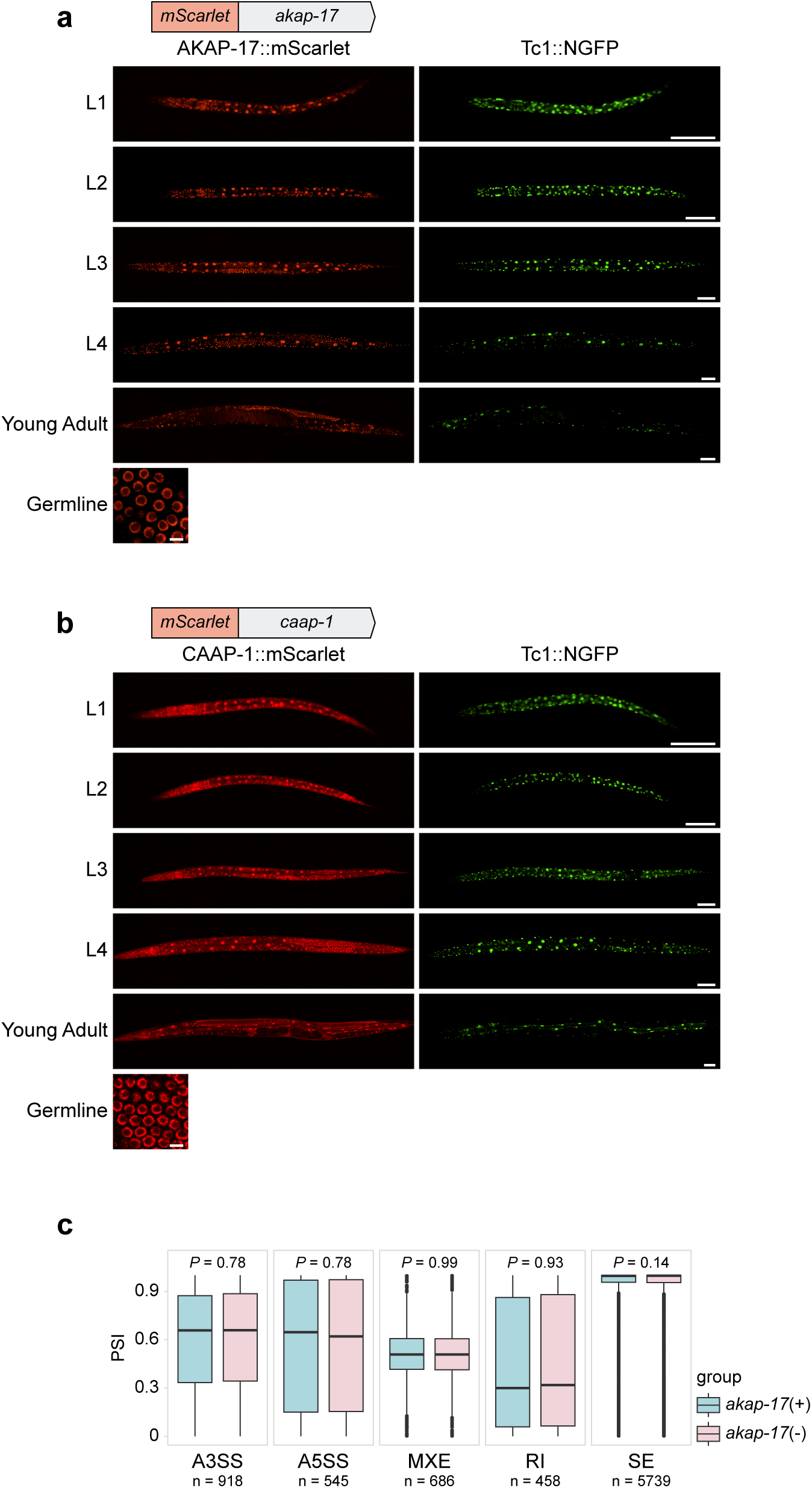
AKAP-17 and CAAP-1 are ubiquitously expressed nuclear proteins, and AKAP-17 is not required for most spliceosomal splicing events. **a-b**. (Top) *mScarlet* was inserted into the N-terminus of the *akap-17* (**a**) or *caap-1* (**b**) genes. (Bottom) Spinning disc confocal images of mScarlet::AKAP-17 (**a**) or mScarlet::CAAP-1 (**b**) in *C.elegans* of the indicated developmental stages. Scale bar = 50 μm for larval stage worms; scale bar = 10 μm for (60 x) images of germ cells. **c**. PSI analysis for five types of alternative splicing events analyzed by rMATS from Illumina-based RNA sequencing data of rRNA-depleted total RNA from animals of the indicated genotypes. n, number of alternative splicing events detected. A3SS, alternative 3’ splice sites; A5SS, alternative 5’ splice sites; MXE, mutually exclusive exons; RI, retained introns; SE, skipped exons. Box and Whisker plots represent the median extends from the 25th to 75th percentiles (box) or min and max values (whiskers) with outliers (outside of 1.5 × interquartile range) shown. The Wilcoxon rank-sum test was performed to assess statistical differences, and all *P*-values are shown.

**Extended Data Figure 9.**
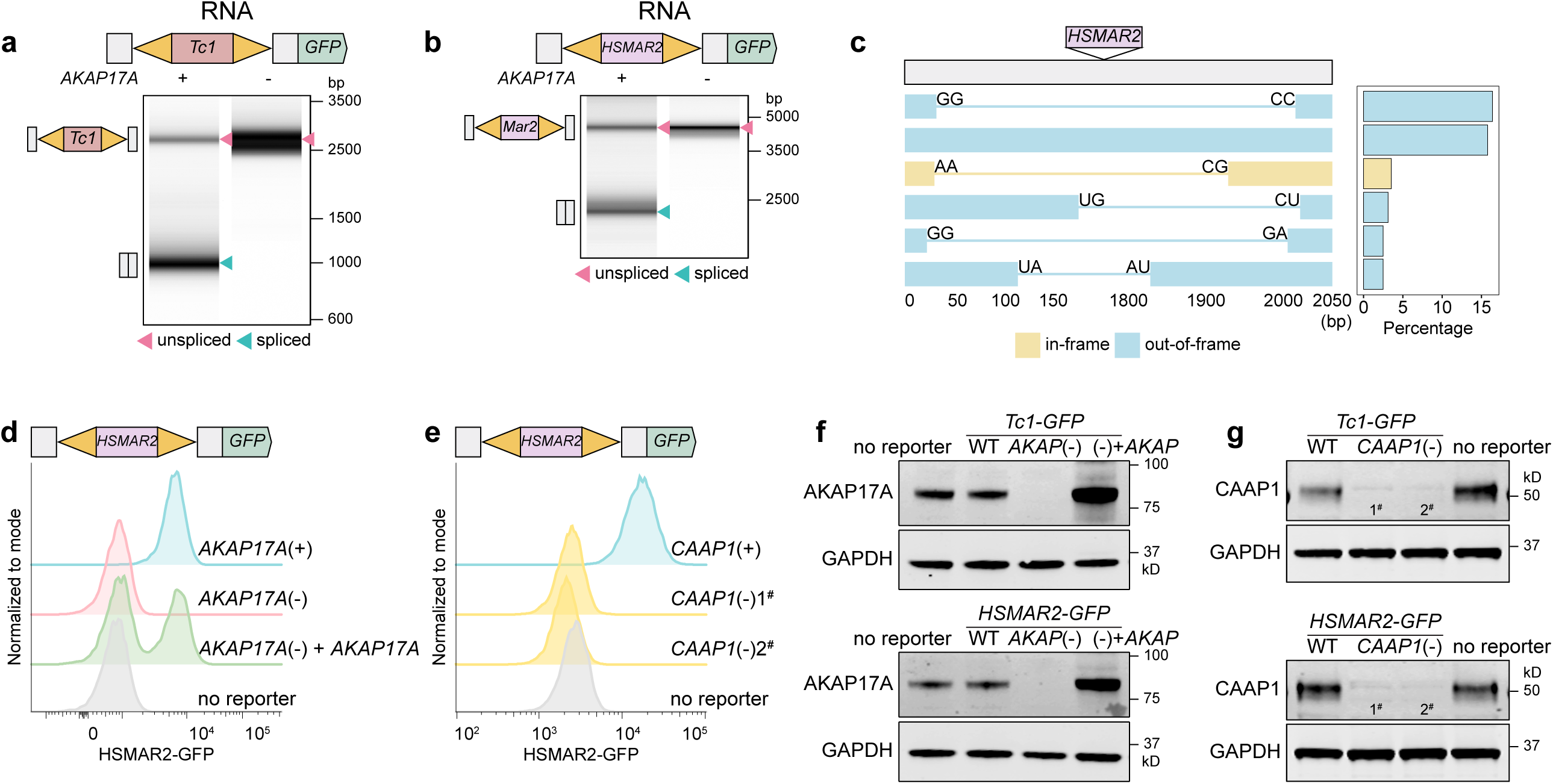
The SOS splicing system is conserved in human cells. **a-b.** Tapestation analysis of amplicons generated from RNA extracted from HEK293T cells of the indicated genotypes, transfected with *Tc1-GFP* (**a**) or *HSMAR2-GFP* (**b**) SOS splicing reporter plasmid. The data show that AKAP17A is needed for SOS splicing of transposons from their respective mRNAs in human cells. Unspliced and spliced amplicons are indicated with arrows **c**. SOS splicing isoforms detected after nanopore-based sequencing of the *HSMAR2-GFP* mRNA, ranked by abundance. Isoforms representing > 2% of total reads are shown. **d-e.** Flow cytometry of HEK293T cells harboring the chromosomally integrated *HSMAR2-GFP* SOS splicing “knock-in” reporter gene. (**d**) *AKAP17A*(+) or (−) cells, and *AKAP17A*(−) cells complemented with *AKAP17A*(+), are shown. (**e**) *CAAP1*(+) cells and two independently derived clones (1# and 2#) of *CAAP1*(−) HEK293T cells are shown. The data show that SOS splicing of the *HSMAR2-GFP* mRNA requires AKAP17A and CAAP1. **f.** Western blot analysis showing that *AKAP17A*(−) cells lack AKAP17A. And that lentiviral-based complementation with wild-type *AKAP17A*(+) was successful. GAPDH serves as a loading control. **g.** Western blot analysis showing that *CAAP1*(−) cells lack CAAP1. GAPDH serves as a loading control.

**Extended Data Figure 10.**
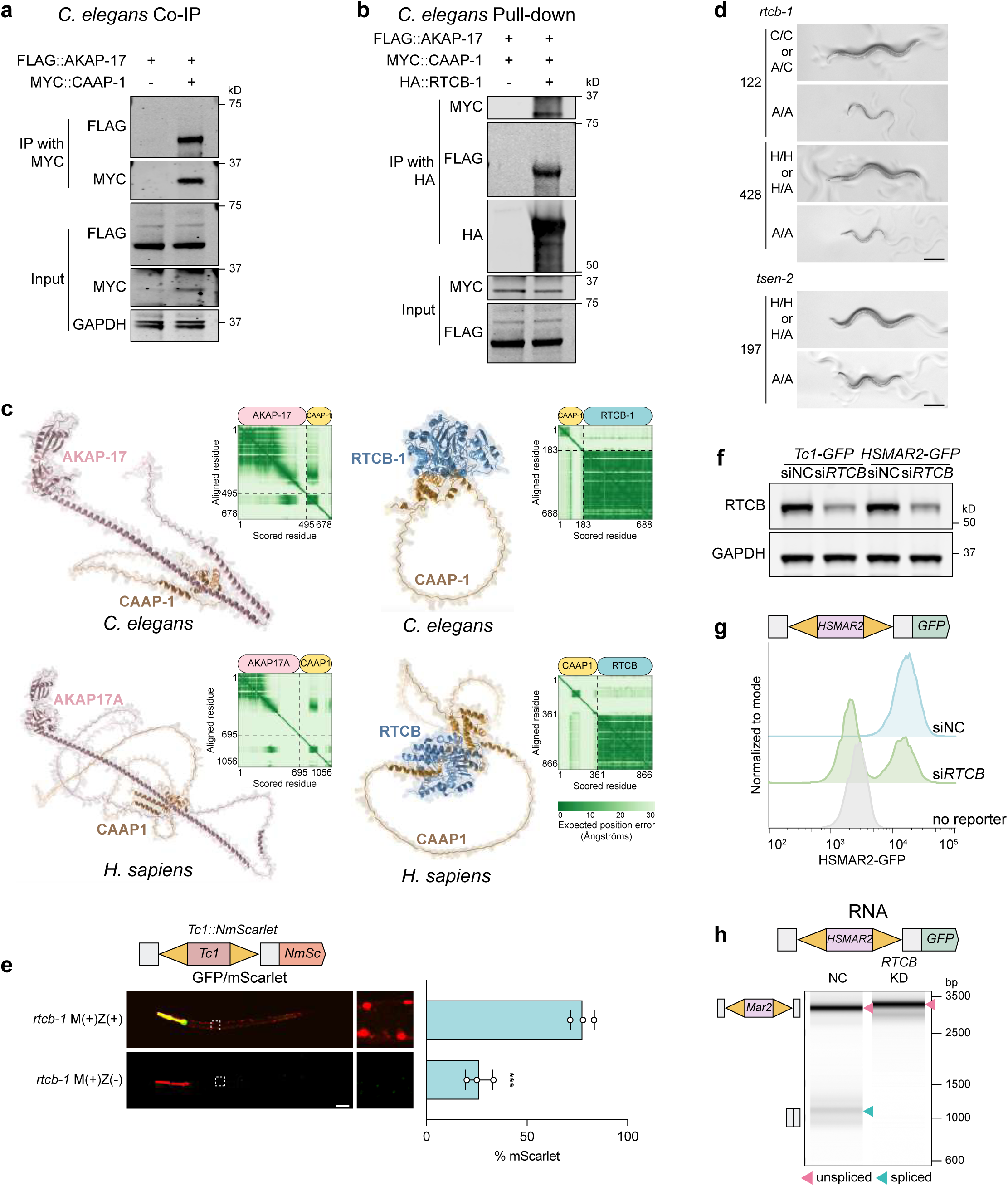
The RNA ligase RTCB is required for SOS splicing. **a.** Co-immunoprecipitation (Co-IP) assay showing interactions between *C. elegans* AKAP-17 and CAAP-1. GAPDH serves as a loading control. **b.** Pull-down experiment demonstrating a physical interaction between *C. elegans* AKAP-17, CAAP-1 and RTCB-1. *C. elegans* HA::RTCB-1 protein was synthesized using PURExpress^®^ *In Vitro* Protein Synthesis Kit, and incubated with lysates generated from animals expressing FLAG::AKAP-17 and MYC::CAAP-1. HA::RTCB-1 was IP’ed with Pierce™ anti-HA magnetic beads and HA::RTCB-1 co-precipitating proteins were detected by Western blot using the indicated antibodies. **c.** AlphaFold Multimer predictions. (Left panel, upper) *C. elegans* AKAP-17 (pink) and CAAP-1 (gold) (ipTM = 0.47). (Left panel, bottom) *H. sapiens* AKAP17A (pink) and CAAP1 (gold) (ipTM = 0.36). (Right panel, upper) *C. elegans* RTCB-1 (blue) and CAAP-1 (gold) (ipTM = 0.3). (Right panel, bottom) *H. sapiens* RTCB (blue) and CAAP1 (gold) (ipTM = 0.23). PAE matrix for each prediction is shown. Dashed black lines indicate chain boundaries. **d.** DIC images of wild-type (N2), *rtcb-1(C122A)*, *rtcb-1(H428A)* (top), and *tsen-2(H197A)* animals (bottom) worms, after 65 hours of development at 20 °C. The data show that *rtcb-1* and *tsen-2* mutant animals arrest development at the L3 or L4 stage, respectively. Scale bar = 200 μm. Genotypes were established by PCR analysis of arrested and non-arrested animals. **e.** *rtcb-1(gk451)* contains a 370 bp deletion spanning the promoter and the first exon of the *rtcb-1* locus. *Tc1::NmScarlet* SOS splicing reporter was injected into *rtcb-1(gk451)*/*hT2 [bli-4(e937) let-?(q782) qIs48]* P0 adult animals, and the percentage of F1 progeny expressing mScarlet signal was quantified. *qIs48* is an insertion of ccEx9747 with markers: MYO-2::GFP expressed brightly in the pharynx throughout development, PES-10::GFP expressed in embryos, and a gut promoter driving GFP in the intestine. GFP marks the wild-type copy of *rtcb-1.* Therefore, lack of GFP expression in progeny indicates animals are *rtcb-1(gk451)* and, therefore, animals not expressing GFP are (*rtcb-1* M(+)Z(−)). Representative images of NmScarlet expression in animals expressing GFP ((*rtcb-1* M(+)Z(+)) and not expressing GFP (*rtcb-1* M(+)Z(−)) are shown. The data show that animals homozygous for a deletion allele of *rtcb-1* are defective for SOS splicing of *Tc1::NmScarlet* SOS splicing reporter. Data are presented as mean ± SD with all data points shown. *N* = 3 biologically independent experiments. The two-tailed unpaired student’s *t*-test. \*\*\**P* < 0.001. Scale bar = 50 μm. **f.** Western blot analysis showing *RTCB* knockdown following si*RTCB* transfection into HEK293T cells. GAPDH serves as a loading control. **g.** Flow cytometry analysis detecting GFP in HEK293T *HSMAR2-GFP* reporter knock-in cells treated with esiRNA targeting *RTCB* (si*RTCB*) or Renilla Luciferase (siNC). **h.** Tapestation analysis of amplicons generated from RNA extracted from HEK293T cells treated with esiRNA targeting RTCB (KD) or Renilla Luciferase (NC) and co-transfected with *HSMAR2-GFP* reporter plasmid. Unspliced and spliced amplicons are indicated with arrows.

